# Essential roles of plexin-B3^+^ oligodendrocyte precursor cells in the pathogenesis of Alzheimer’s disease

**DOI:** 10.1101/2020.03.30.015297

**Authors:** Naomi Nihonmatsu-Kikuchi, Xiu-Jun Yu, Yoshiki Matsuda, Nobuyuki Ozawa, Taeko Ito, Kazuhito Satou, Tadashi Kaname, Akihiko Takashima, Shuta Toru, Katsuiku Hirokawa, Masato Hasegawa, Toshiki Uchihara, Yoshitaka Tatebayashi

## Abstract

The roles played by oligodendrocyte (OL) lineage cells, the largest glial population in the adult CNS, in the pathogenesis of Alzheimer’s disease (AD) remain elusive. Here, we show a newly developed culture method for adult OL progenitor cells (aOPCs) and identify novel plexin-B3-expressing (plexin-B3^+^) aOPCs as potential amyloid β peptides (Aβ)-secreting cells. Fibroblast growth factor 2 (FGF2) promotes the survival and proliferation of aOPCs in a serum-free defined medium. Although the whole expression profiles of the expanded aOPCs closely resemble those of in vivo OPCs, we found a subpopulation (up to 5%) of plexin-B3^+^/olig2^+^ aOPCs in the cultures growing in FGF2. FGF2 withdrawal decreased NG2^+^, but increased plexin-B3^+^ aOPCs with increased APP expression, Aβ1-40, −42 secretions and Aβ1-42/total Aβ ratios in association with cored senile plaque-like morphological changes. In vivo, plexin-B3^+^ aOPCs are distributed throughout the adult brain, although less densely so than NG2^+^ aOPCs. Spreading depolarization, a type of brain injury, induced unique delayed cortical plexin-B3^+^ aOPC gliosis in the ipsilateral, but not in the contralateral, remote cortex. In AD brains, virtually all senile plaques in the cortex were immunostained with plexin-B3 antibodies and the levels of cortical plexin-B3 expression increased significantly in the Salcosyl-soluble fractions. These findings suggest that plexin-B3^+^ aOPCs play important roles in the pathogenesis of AD most likely as a natural Aβ source.

## Introduction

The amyloid hypothesis for Alzheimer’s disease (AD) posits a neuron-centric, linear cascade initiated by the abnormal production of longer forms of amyloid β peptides (Aβ), especially Aβ1-42, from amyloid precursor protein (APP), leading progressively to tau pathology, synaptic dysfunction, inflammation, neuronal loss, and, ultimately, dementia^1^. Activated astrocytes and microglia are commonly found as glial nests around senile plaques (SPs) composed of Aβs^2^. Such reactive gliosis occurs as a multicellular neuroinflammatory response to Aβ accumulation and is believed to contribute to the clearance and removal of extracellular Aβs. While much attention has, therefore, been paid to these cells^1^, the roles played by other major glia, such as oligodendrocyte (OL) lineage cells, in the pathogenesis of AD remain largely unknown.

OL lineage cells constitute ∼75% of the neuroglial cells in the neocortex and are thus the largest group of non-neuronal cells in the adult human brain^3^. While mature OLs produce myelin and facilitate neuronal transmission, the roles played by adult OL progenitor cells (aOPCs), which are characterized by the expression of the platelet-derived growth factor receptor α subunit (PDGFRα) and the NG2 proteoglycan^4, 5^, are still unknown. These aOPCs, which descend from OPCs in the perinatal CNS^6^, are distributed throughout the adult brain, providing up to 5 – 10% of the adult CNS cells^7^. Although they generally continue to proliferate, some generate myelinating OLs in the gray and white matter even during adulthood^7, 8^. Furthermore, in the human prefrontal and entorhinal cortex, myelination in the gray matter continues throughout life, peaking around the 5^th^ - 6^th^ decade^9, 10^.

Braak & Braak found a link between the vulnerability of neurons and the cortical (gray matter) myelination due to the fact that the spreading of neurofibrillary tangles recapitulates the developing pattern of cortical myelination during adulthood in reverse order^11^. This neuropathological finding, though clearly demonstrating the close relationship between adult gray matter OL lineage cells and the AD pathogenesis, has been underestimated, partly due to the lack of appropriate markers and/or in vitro systems to investigate such relationships in humans and rodents. Most recently, Mathys and colleagues illuminate the brain transcript of AD at single-cell resolution and found unexpectedly that, at the early stage of AD, myelination and axonal integrity and repair may be essentially involved in the pathogenesis^12^. However, little is known about how OL lineage cells change with healthy aging and what their role is in the initiation and progression of disease^13^.

Although CNS cell purification and culture from embryonic or postnatal (up to approximately P20) rodent brains are possible using several different protocols, those from adult (> 2 months old) brains remain more challenging and often result in no or low yield(s), except for a few cell types (i.e., adult microglia and hippocampal neural stem/progenitor cells (NSPCs) that reside within a highly specific stem cell niche in the dentate gyrus^14–17^. Culturing OL lineage cells from adult brains is rather complex. While OPCs from postnatal ∼ adult rodent optic nerves can be purified by using a combination of sequential immunopannings^18^, advancing age, especially after P50, renders any attempt to culture them in vitro increasingly difficult, and in fact nearly impossible^19^. Utilizing surgically resected adult human brains, Antel’s group has established a method for purifying and culturing OLs (or OPCs) that can survive for up to 4 weeks in vitro^20, 21^. However, the use of serum limits their precise functional analysis and purity.

In the present study, we developed a novel method to purify aOPCs from the adult rat brain (> 2 months old) and to culture them for up to several months in a defined medium. This allowed us to identify plexin-B3 as a novel uncharacterized aOPC marker. We also found that plexin-B3 is a unique delayed marker for cortical gliosis following brain injuries. Furthermore, we show that plexin-B3^+^ aOPCs probably play roles in the pathogenesis of AD, most likely as natural Aβ (especially Aβ1-42)-secreting cells.

## Results

### Purification and culture of aOPCs from adult rat brain

Density gradients have often been used to purify adult microglia and hippocampal neural stem/progenitor cells from adult rodent brains^15–17^. This allows less buoyant cells to be separated on density gradients spanning 1.030 – 1.065 g/ml^14–16^ in the case of microglia, and 1.065–1.074 g/ml in the case of NSPCs^17^. In these purification procedures, the fractions with much greater buoyancy (<1.030 g/ml) were always discarded because they were regarded as merely accumulated debris^14, 15, 17^. However, we found that many unidentified cells were present in this more buoyant fraction (1.029 g/ml) (Fig. 1a).

**Fig. 1.**
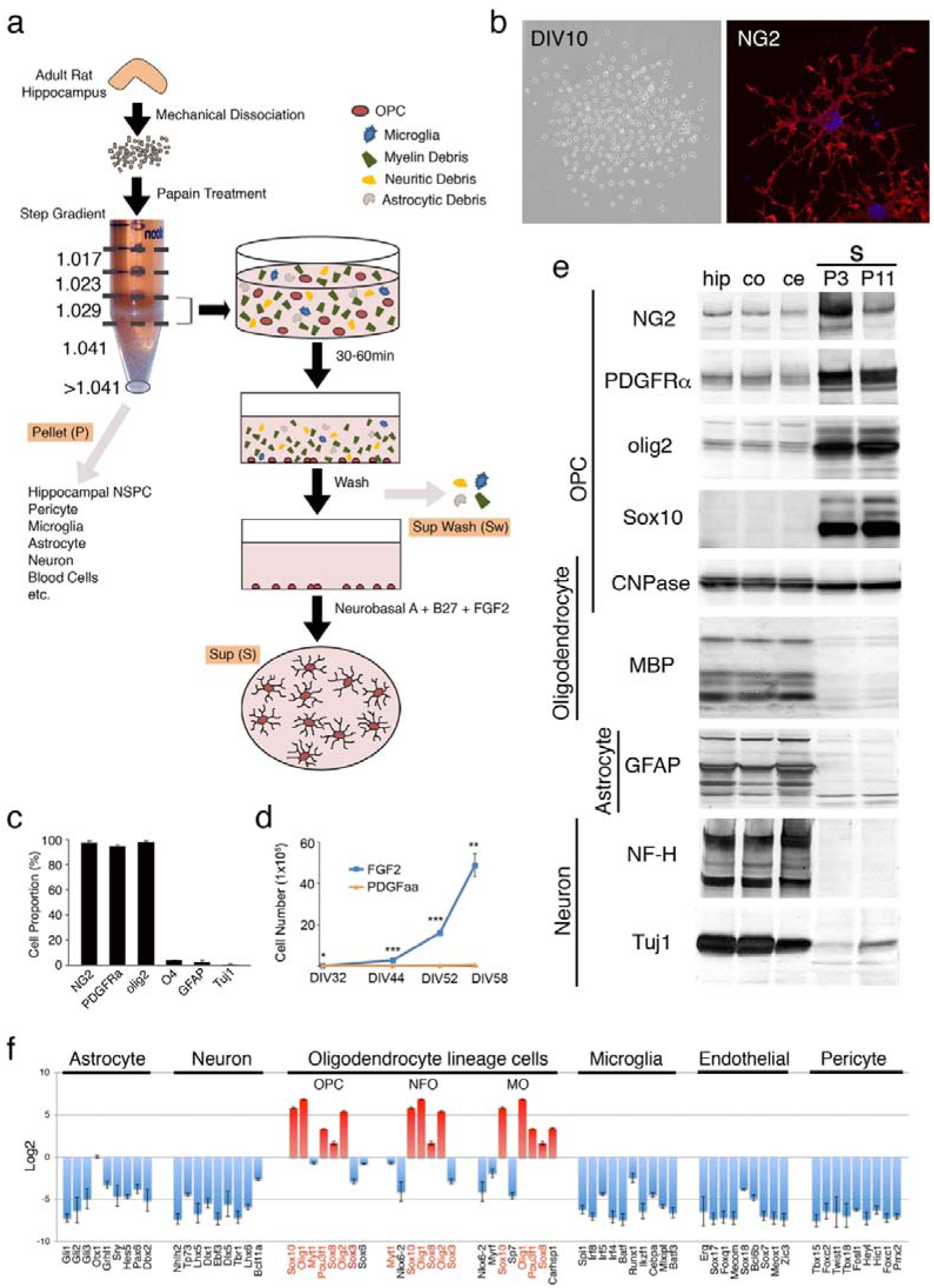
Purification and culture of aOPCs. (a) Culture strategy. NSPC: neural stem or precursor cells. (b) Left: Phase contrast images of primary cultures at 10 days in vitro (DIV10). Right: Morphology of a NG2^+^ cell. (c) Proportions of marker-positive cells (mean±SD) at DIV 5. (d) Growth effect of FGF2 compared to PDGFaa after passage 1 (\*\*\**P* < 0.0001, \*\**P* < 0.001, \**P* < 0.005). (e) Protein expression profiles of cultured aOPC. P3: aOPCs at passage 3; P11: passage 11; CNPase: 2’,3’-Cyclic-nucleotide 3’-phosphodiesterase; PLP: Proteolipid protein; MBP: Myelin basic protein; MAG: Myelin-associated glycoprotein; GFAP: Glial fibrillary acidic protein; NF-H: neurofilament H; Tuj1: neuron-specific class III β-tubulin; hip: adult rat hippocampus; co: cortex; ce: cerebellum. (f) Transcription factor gene expression profiles of microarray of cultured aOPCs. The cell-specific transcription factor genes were introduced from Zhang et al. (2014) (Zhang et al., 2014). Magenta: highly expressed genes; Blue: downregulated genes; Red genes: overlapped genes in oligodendrocyte lineage cells. See also Supplementary Table 4.

Preliminary immunolabeling studies revealed that most cells in this fraction were positive for olig2, a transcriptional factor and a marker specific for OL lineage cells, while a smaller population was positive for Iba1, a marker for microglia. To further purify olig2^+^ cells, we overlaid these fractions on poly-d-lysine-coated dishes for 30 – 60 min and then gently washed out the suspension. Because olig2^+^ cells more quickly and tightly adhered to the dishes, most likely due to their membrane charges, most microglias and debris were removed to the wash suspensions (Sw) (Fig. 1a).

In Neurobasal A/B27 supplemented with FGF2, some of the attached cells began to proliferate (Fig. 1b). After 5 DIV, more than 90% of the cells were co-immunolabeled with antibodies to NG2, PDGFRα, and olig2 (Fig. 1c).

Only in the primary cultures was a mitogenic effect also found in platelet-derived growth factor aa (PDGFaa), although not in epidermal growth factor, nerve growth factor, neurotrophin-3, or in ciliary neurotrophic factor. The effects of PDGFaa on cell morphology and migration were different from those of FGF2 (Supplementary Fig. 1). After the first passage, however, only FGF2 showed continuously increasing numbers of aOPCs (Fig. 1d). Within a few passages in FGF2, the cultures became more homogeneous; in fact, nearly 100 % of the cells became olig2^+^ and more than 95 % of those NG2^+^.

Western blot (WB) analysis revealed strong expression of OPC markers including NG2, PDGFRα, olig2, and Sox10 (Fig. 1e). Markers for mature OLs, astrocytes, or neurons remained undetected or only detectable at negligible levels (Fig. 1e). Microarray (Fig. 1f) and RNA-seq (Supplementary Fig. 4A) analyses further confirmed that they expressed OL lineage (especially OPC)-specific transcription factor genes such as Sox10, Olig1, Pou3f1, Sox8, and Olig2^22^. Comparison of the microarray (Supplementary Fig. 2) and RAN-seq (Supplementary Fig. 3) profiles of cultured aOPCs with two single-cell transcriptome databases of adult mouse CNS cells by RNA-seq (Top 40 (Zhang et al., 2014) ^22^ and Top 50 (Wu, Pan et al., 2017)^23^) further confirmed the relatively specific expression of the OPC genes in the cultured cells. Taken together, these data suggest not only that OL lineage cells, especially aOPCs, can be successfully isolated and cultured from the adult rat hippocampus, but also that FGF2 can effectively maintain aOPC properties for at least several months in vitro.

### Plexin-B3^+^ aOPCs in vitro

In our analyses of microarray and RNA-seq, we found that a transmembrane protein, plexin-B3^24^, was highly expressed in cultured aOPCs. This was further confirmed in the WB analysis (Fig. 2a). Although plexin-family members are generally expressed in neuronal cells, Zhang et al. previously reported in an transcriptome database that plexin-B3 gene was enriched in OL lineage cells, especially in newly formed OLs (NFO)^22^ (Zhang et al., 2014). Immunocytochemical studies further revealed that plexin-B3^+^ cells comprised less than 5% of total cells and all intensely olig2^+^ (Fig. 2b & c). Plexin-B3^+^/olig2^+^ cells are generally NG2^−^ or very weakly NG2^+^ (Fig. 2b). Moreover, BrdU incorporation studies revealed that they even proliferate in culture (Fig. 2e), suggesting that plexin-B3 is most likely a novel aOPC marker.

**Fig. 2.**
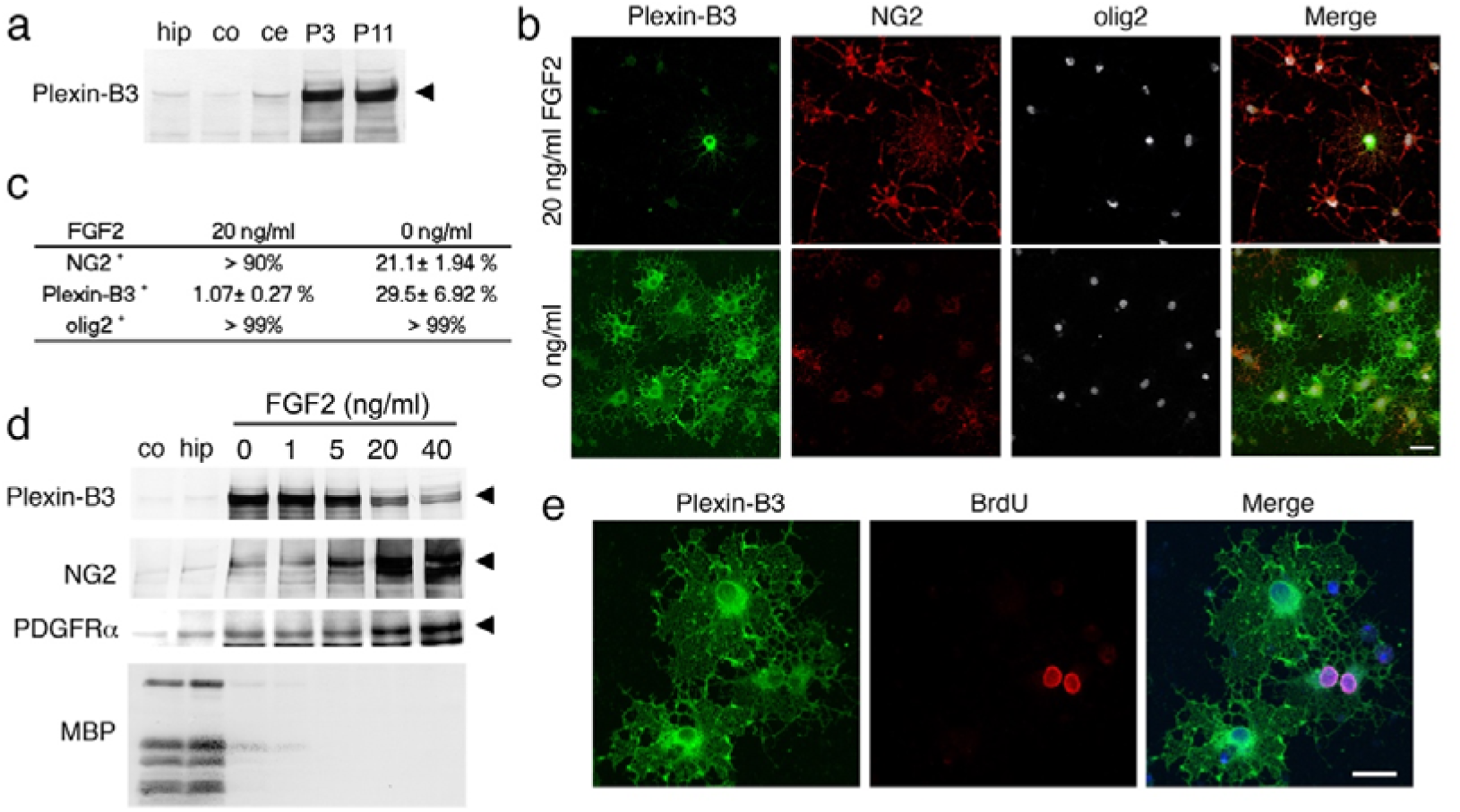
Characterization of plexin-B3^+^ aOPCs in vitro. (a) Plexin-B3 expression of cultured aOPCs. P3: aOPCs at passage 3; P11: passage 11; hip: adult rat hippocampus; co: cortex; ce: cerebellum. (b) Effects of FGF2 (0 or 20 ng/ml) on the expression of plexin-B3, NG2, and olig2. Scale bar: 20 μm. (c) Quantitative cell counting of cultured aOPCs. (d) Effects of FGF2 on the levels of plexin-B3, NG2, PDGFRα, and MBP. (e) BrdU incorporation of plexin-B3^+^ aOPCs. Scale bar: 20 μm.

Since mitogen withdrawal induces OL differentiation of cultured perinatal OPCs^28^, the effect of FGF2 withdrawal on the expression of the aOPC marker was studied. FGF2 withdrawal of cultured aOPCs for 5 days dramatically increased the proportion of plexin-B3^+^ aOPCs (29.5 ± 6.9 %) and decreased the proportions of NG2^+^ aOPCs (27.9 ± 3.4 %) without changing the proportions of olig2^+^ cells (> 99 %) (Fig. 2b, c, & Supplementary Fig. 4D, see FGF 0 ng/ml). FGF2 withdrawal also induced uniquely ramified or, occasionally, cored SP-like morphological changes (Fig. 2b & Fig. 3e). WB analysis further confirmed that FGF2 withdrawal or the FGF2 inhibitor PD173074, dose-dependently increased plexin-B3 (Fig. 2d & Supplementary Fig. 4E), whereas FGF2 withdrawal decreased NG2 and PDGFRα levels (Fig. 2d).

**Fig. 3.**
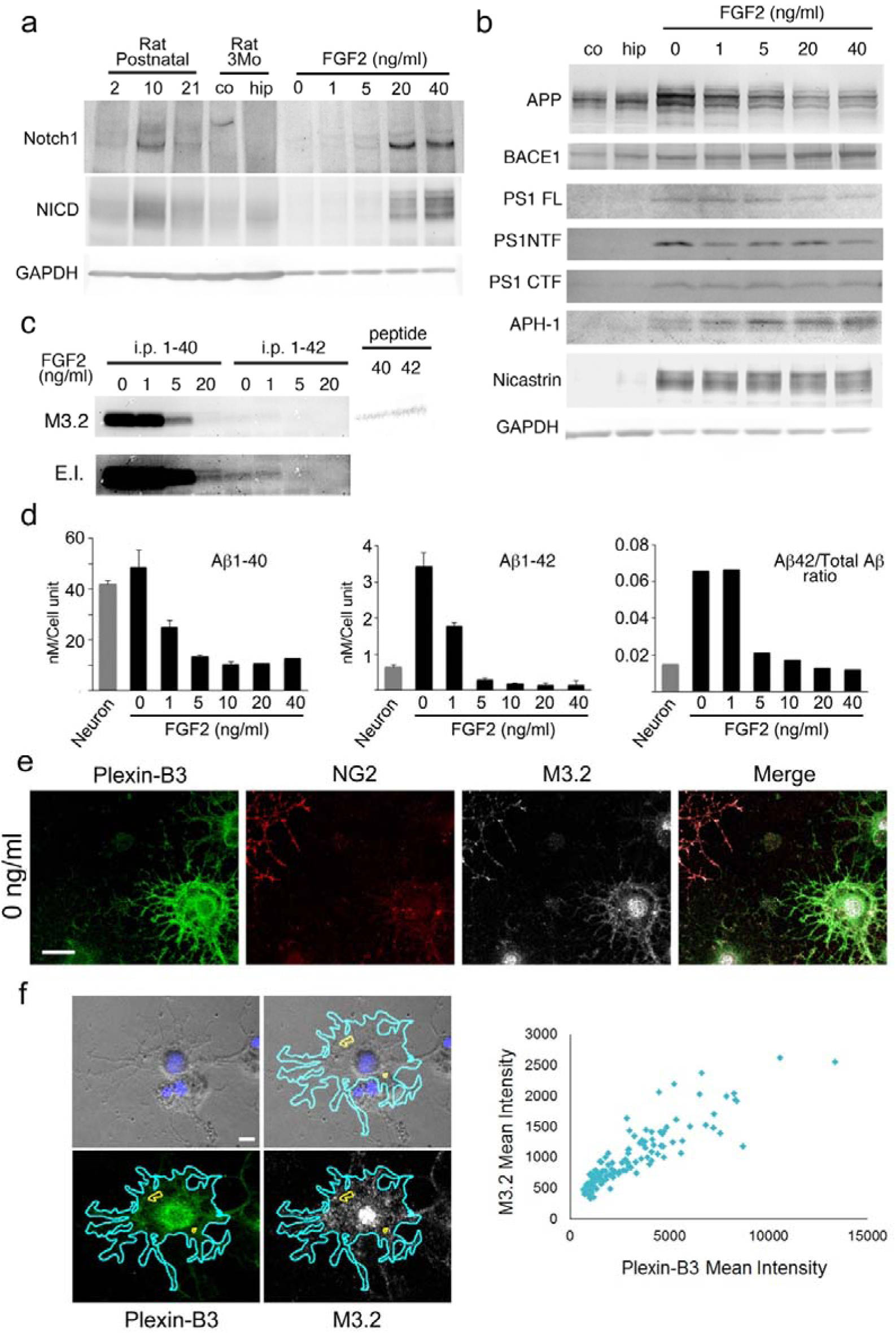
Notch and APP processing in cultured aOPCs. (a) Expression of Notch1 and its intracellular signal mediator NICD in cultured aOPCs. Their expression levels were also compared with those from rat brains at postnatal day 2, 10, and 21 and adult rat cortex (co) and hippocampus (hip). Aliquots of rat brains (100 μg) and cultured aOPCs (15 μg) were separated by SDS-PAGE and subjected to WB analysis with Notch1, NICD, and GAPDH. GAPDH: Glyceraldehyde-3-phosphate dehydrogenase. (b) Effects of FGF2 on the levels of APP and β- and γ-secretases. BACE1: β-site amyloid precursor cleaving enzyme 1 (β-secretase); PS1 FL: presenilin 1 full length; PS1 NTF: PS1 N-terminal fragment; PS1 CTF: PS1 C-terminal fragment; APH-1: anterior pharynx-defective 1. (c) WB analysis (M3.2) of immunoprecipitated (i.p.) Aβs. M3.2: monoclonal antibody for rat Aβs (Aβ10−15). peptide 40: human Aβ40 detected by monoclonal antibody 4G8; peptide 42: human Aβ42; E.I.: enhanced image. (d) ELISA for rodent Aβx-40 and x-42. Amounts of secreted Aβx-40 and x-42 from cultured fetal rat hippocampal neurons (3 weeks in vitro) are also shown. (e) Immunocytochemistry of a typical cored-SP-like plexin-B3^+^ aOPC cultured in 0 ng/ml FGF2 for 5 days. Note that plexin-B3^+^ aOPC is also M3.2 positive. Scale bar: 15 μm. (f) Quantitative analysis of plexin-B3 and M3.2 (full length APP or intracellular Aβ) immunoreactivities. An example of region of interest for a single cell is shown in the left panels. Note that, as plexin-B3 immunoreactivities increased, M3.2 immunoreactivities increased (right). Mean Intensity: the integrated total value of the signal intensities adjusted by the single cell area.

We found that the 5-day FGF2 withdrawal did not clearly change the transcription factor gene expression profiles of cultured aOPCs (Supplementary Fig. 4A). In more detailed analyses using the lists of OL lineage-specific genes (the Top 40 genes of OPC, NFO, and MO by Zhang et al. ^22^ and the Top 40 genes of OPC and oligodendrocyte by Wu et al.^23^), we found that the 5-day FGF2 withdrawal increased several NFO- or MO- (or oligodendrocyte-)specific gene expression profiles (Supplementary Fig. 4B & 4C), although many OPC genes (such as Gpr17) were still highly expressed in cultured aOPCs.

Interestingly, the protein levels of myelin basic protein (MBP), an essential marker for mature OLs and myelin, increased only faintly (Fig. 2d & Supplementary Fig. 4F), although the mRNA levels became extremely high after 5-day FGF2 withdrawal (Supplementary Fig. 4B & 4F). This phenomenon may represent a type of post-translational repression of MBP protein, since its protein synthesis is strictly regulated by the correct temporal and spatial transport of mRNA and its localized translation into the protein^26, 27^.

Taken together, these results indicate that FGF2 withdrawal does initiate OL differentiation of cultured aOPCs, albeit incompletely in vitro, and that plexin-B3 is most likely an uncharacterized late OPC marker.

### Notch and APP processing in plexin-B3^+^ aOPCs in vitro

Since notch signaling is known to inhibit OL differentiation and myelination^28^ of cultured perinatal OPCs, we next studied the effects of FGF2 withdrawal on notch signaling in cultured aOPCs. WB analysis revealed that FGF2 dose-dependently increased both of the notch1 and notch1 intracellular domains (NICDs) generated by γ-secretase (Fig. 3a), suggesting that cultured aOPCs do possess regulatory systems for notch and γ-secretase. FGF2 dose-dependently increased notch signaling, supporting the notion that notch signaling might inhibit OL differentiation of cultured aOPCs.

We then investigated the processing of APP, another major substrate for γ-secretase, in vitro. APP was expressed in cultured aOPCs and FGF2 withdrawal increased its expression (Fig. 3b & Supplementary Fig. 4B). The levels of APP protein in cultured aOPCs with 0 or 1 ng/ml FGF2 were much higher than those of adult rat hippocampus and cortex tissues (Fig. 3b), suggesting that cultured aOPCs express high levels of APP in vitro depending on the culture conditions. The β-secretase BACE1^29^ and the components of γ-secretase, including presenilin-1 (PS1)^30, 31^, nicastrin^32^, and Aph-1^33, 34^, were also highly expressed in cultured aOPCs (Fig. 3b). Interestingly, the band patterns of Aph-1 and nicastrin changed depending on the concentration of FGF2 (Fig. 3b).

We next studied the secretion of Aβs. Aβ1-40 or Aβ1-42 peptides were immunoprecipitated from the conditioned medium with antibodies specific for the carboxyl (C-) terminal of Aβ1-40 or Aβ1-42, respectively, and were analyzed by WB with an M3.2 antibody that recognizes rodent Aβ10-15. As a control, secreted APPs (sAPPs) were also immunoprecipitated with the monoclonal antibody 22c11 (Supplementary Fig. 5A & B). Unlike NICD generation, the levels of secreted Aβ1-40 and −42 were most abundant in 0 ng/ml FGF2 (Fig. 3c), even after adjusting for the levels of sAPPs (Supplementary Fig. 5A & B), suggesting that γ-secretase also cleaved APP in cultured aOPCs, albeit in a manner completely opposite to that of Notch1 processing. The underlying mechanism of this unique observation in cultured aOPCs is currently unknown, however, changes in the substrate levels (Fig. 3a Notch1 and 3b APP) or potentially the substrate preference of γ-secretase as implicated from the bands shifts of APH-1 and nicastrin depending on the concentrations of FGF2 (Fig. 3b) might be involved.

To more quantitatively measure the levels of Aβ in the medium, we performed ELISAs for Aβx-40 or Aβx-42. An LDH assay was also performed in parallel to adjust cell numbers. To compare Aβ levels between cultured aOPCs and neurons, primary fetal rat hippocampal neurons were also cultured (3 weeks in vitro). We again confirmed that the levels of both Aβx-40 and −42 increased as FGF2 concentrations decreased (Fig. 3d). Notably, aOPCs with 0 or 1 ng/ml FGF2 secreted more Aβx-42 than cultured fetal rat neurons, resulting in approximately 4-fold higher ratios of Aβx-42 to total Aβ in aOPC than in fetal rat neuron cultures (4.39 and 4.45-fold increases, respectively) (Fig. 3d).

Immunocytochemical studies further confirmed that cored SP-like plexin-B3^+^ aOPCs in 0 ng/ml FGF2 expressed APP (Fig. 3e). Quantitative measurements of plexin-B3 and APP (M3.2) levels per single cell further indicated that, as plexin-B3 levels increased, APP levels increased (Fig. 3f). Since almost all of the plexin-B3^+^ cells were olig2^+^ (> 99%, Fig. 2c & Supplementary Fig. 4D), we concluded that aOPCs, especially those expressing plexin-B3, possess the ability to secrete considerable amounts of Aβs naturally (without genetic engineering; i.e., overexpression with point mutations) in vitro.

### Plexin-B3^+^ aOPCs in rat brains

In vivo, plexin-B3^+^ aOPCs were distributed throughout the adult rat brain (Fig. 4a, b & Supplementary Fig. 6 & 7). In the cortex and the hippocampus, anti-plexin-B3 antibodies stained mostly cells with aOPC-like morphologies (Fig. 4a & Supplementary Fig. 7). In the corpus callosum (CC), however, they sometimes stained pre OL-like cells (Fig. 4c & Supplementary Fig. 6), suggesting a possibility that plexin-B3 may remain expressed in the early phase of OL differentiation in vivo.

**Fig. 4.**
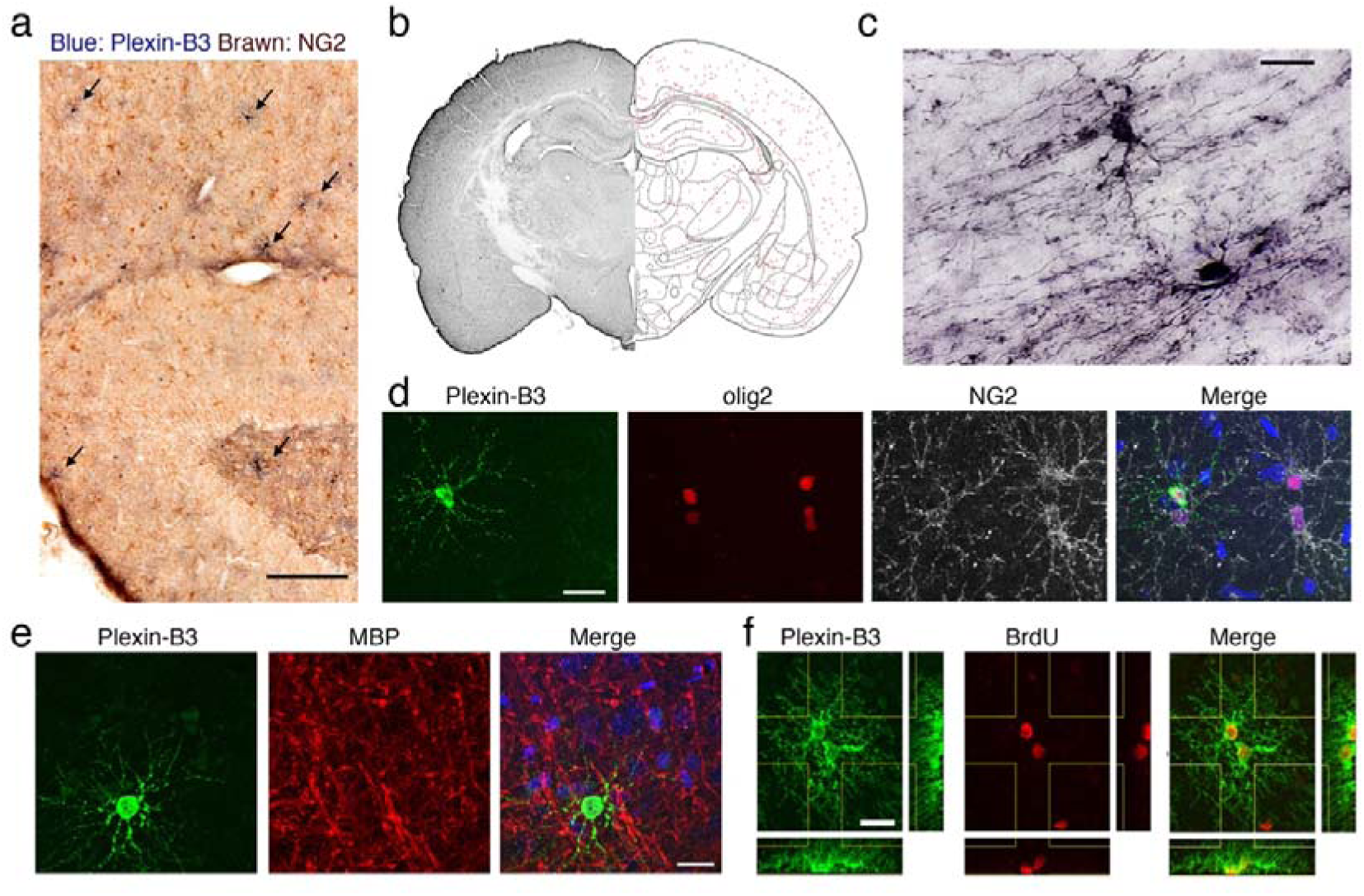
Characterization of plexin-B3^+^ aOPCs in vivo. Immunohistochemistry of plexin-B3 (blue) and NG2 (brown) in the adult rat hippocampus. Arrows: plexin-B3^+^ aOPCs. Scale bar: 200 μm. (b) Distribution of plexin-B3^+^ aOPCs in an adult rat brain section (red dots). (c) Morphologies of plexin-B3^+^ aOPCs in the corpus callosum. Scale bar: 20 μm. (d) Fluorescent immunostaining of plexin-B3, olig2 and NG2 in the cortex. Scale bar: 20 μm. (e) Fluorescent immunostaining of plexin-B3 and MBP in the cortex. In (d) and (e), nuclear stainings (blue) are shown in the merged panels. Scale bar: 20 μm. (f) Fluorescent immunostaining of plexin-B3 and BrdU in the cortex. Scale bar: 20 μm.

They were generally NG2^−^ (Fig. 4d & Supplementary Fig. 6), GFAP- (Supplementary Fig. 7A), Iba1- (Supplementary Fig. 7A), MBP^−^ (Fig. 4e & Supplementary Fig. 6), but they were all intensely olig2^+^ (Fig. 4d & Supplementary Fig. 6), further supporting the contention that they mostly constitute aOPCs (or early phase of pre OL in the CC). The density of plexin-B3^+^ aOPCs in the normal adult rat cortex (∼100/mm^2^) was much lower than in those of olig2^+^ (∼ 1200/mm^2^) or NG2^+^ (∼ 400/mm^2^) aOPCs. Plexin-B3^+^ aOPCs were still proliferative and pairs of plexin-B3^+^/BrdU^+^ aOPCs, as well as occasionally those of plexin-B3^+^/BrdU^+^ and NG2^+^/BrdU^+^ aOPCs, were observed (Fig. 4f & Supplementary Fig. 7B).

### Brain injuries and plexin-B3^+^ aOPCs

It is well known that NG2^+^ aOPCs respond very quickly to brain injuries^35–38^, a major risk factor for AD^42^. To investigate the effects of brain injury on plexin-B3^+^ aOPCs, we first employed the stab wound model. At 2 to 3 days post stab wound, a dramatic response in NG2^+^ aOPCs was first observed in and around the stab lesions with increased NG2 immunoreactivities, as well as hypertrophic morphological changes (Fig. 5a, NG2, 2 Days). Interestingly, no such clear plexin-B3^+^ aOPC response was noted in the same lesions (Fig. 5a, Plexin-B3, 2 Days).

**Fig. 5.**
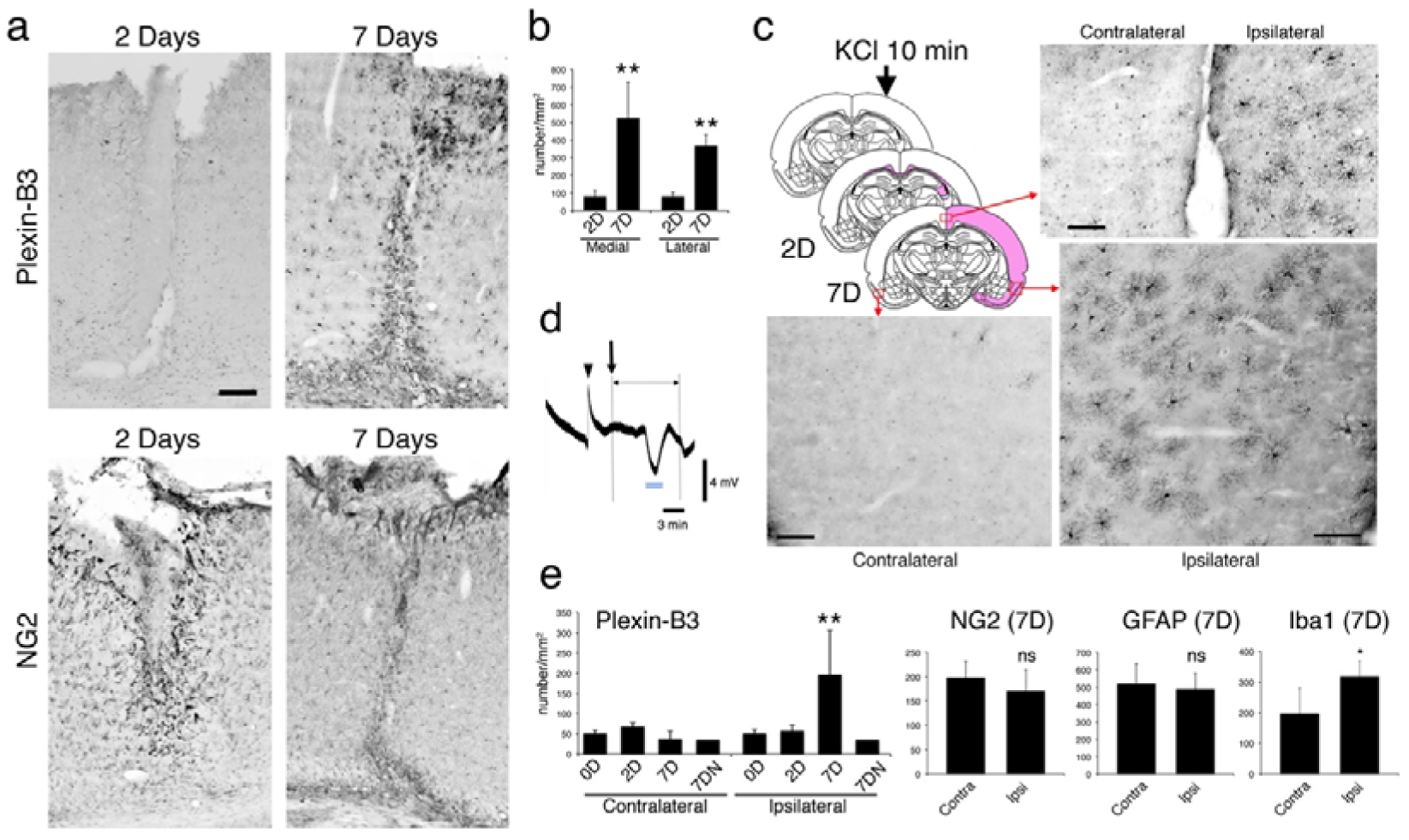
Brain injuries and plexin-B3^+^ aOPCs. (a) Stab wound. Plexin-B3 and NG2 immunohistochemistry at 2 (2D) and 7 (7D) days after the stab insertion are shown. Scale bar: 200 μm. (b) Quantitative cell countings in the defined areas (see Supplementary Fig. 8A) after the injury. \*\**P* < 0.01. (c) Position of KCl injury (arrow), increased plexin-B3 immunoreactive areas (pink coloring) at 2 (2D) and 7 (7D) days, and typical plexin-B3 immunostainings at 7D (from 3 brain areas marked by red rectangles). Scale bar: 100 μm. (d) Representative tracings of direct current potential (millivolts) recorded simultaneously during the 20 min before and after KCl application (arrow). Arrowhead: microelectrode insertion; Blue bars: occurrence of CSD. (e) Quantitative plexin-B3^+^ aOPC numbers in the defined areas (see Supplementary Fig. 8B) at 0D, 2D, and 7D after the KCl application. 7DN: 7D after the NaCl application. Numbers of NG2^+^ aOPCs, GFAP^+^ astrocytes, and Iba1^+^ microglias at 7D were also quantified. \*\**P* < 0.01, \**P* < 0.05.

At 6 to 7 days post stab wound, however, a dramatic increase in the numbers of plexin-B3^+^ aOPCs was observed. On the walls of the stab wound (Fig. 5a), plexin-B3 was intensely expressed in olig2^+^ aOPCs, which became hypertrophic and formed glial scars (Fig. 5a). In the gray matter near these glial scars, both laterally and medially, densities of plexin-B3^+^ aOPCs had significantly increased (Fig. 5b & Supplementary Fig. 8A).

Next, we employed the KCl injury model. Topical application of 3 M KCl for 10 minutes induced cortical spreading depression (CSD) within the ipsilateral cortex (Fig. 5c & d). CSD is a self-propagating wave of cellular depolarization that has been implicated as a fundamental mechanism of progressive cortical injury observed in stroke and head trauma^40, 42^. At 2 to 3 days post-KCl application, increased plexin-B3 immunoreactivity was found only within the CC, just beneath the necrotic lesions of the KCl (Fig. 5c, pink-colored area in 2D); however, these white matter reactions decreased to normal levels within 6 to 7 days.

At 6 to 7 days post-KCl application, however, plexin-B3^+^ aOPCs increased not only around the necrotic cortical lesions but also in the remote ipsilateral cortex (Fig. 5c & e). Slight microgliosis, though not NG2^+^ aOPC gliosis or astrocytosis, was observed in the same remote ipsilateral cortical areas in this mild CSD model (Supplementary Fig. 8B). This delayed cortical gliosis was never induced in the contralateral cortex (Fig. 5c & e). NaCl application similarly did not induce such gliosis (Fig. 5e, 7DN).

### Characteristics of plexin-B3 expression in AD brains

Our in vitro and in vivo findings led us to consider a new idea, namely that plexin-B3^+^ aOPCs constitute one of the Aβ-secreting cells in AD. To more directly test this idea, we stained paraffin-embedded human brain sections from patients with AD and normal controls (Supplementary Table 3A). In normal control brains, we could not get any clear plexin-B3 immunoreactive structures in the paraffin sections. In AD brains, however, we found that plexin-B3 antibodies stained almost all SPs, including cored ones (Fig. 6a & b, Supplementary Fig. 9, 10 & 11). These plexin-B3^+^ structures were specific (Fig. 6c & d, Supplementary Fig. 11A), and co-immunolabeled with antibodies for total Aβ (4G8 and Aβ1-42 (Fig. 6e∼h & Supplementary Fig. 10). Interestingly, Aβ^+^ areas of cored SPs were always slightly larger than the corresponding plexin-B3^+^ cell body areas (Fig. 6e∼h, merge, see also Supplementary Fig. 10), indicating the extracellular precipitation of Aβs. We also confirmed that these plexin-B3 SPs were closely associated with, but clearly distinct from, microglia and astrocytes (Fig. 6i∼l).

**Fig. 6.**
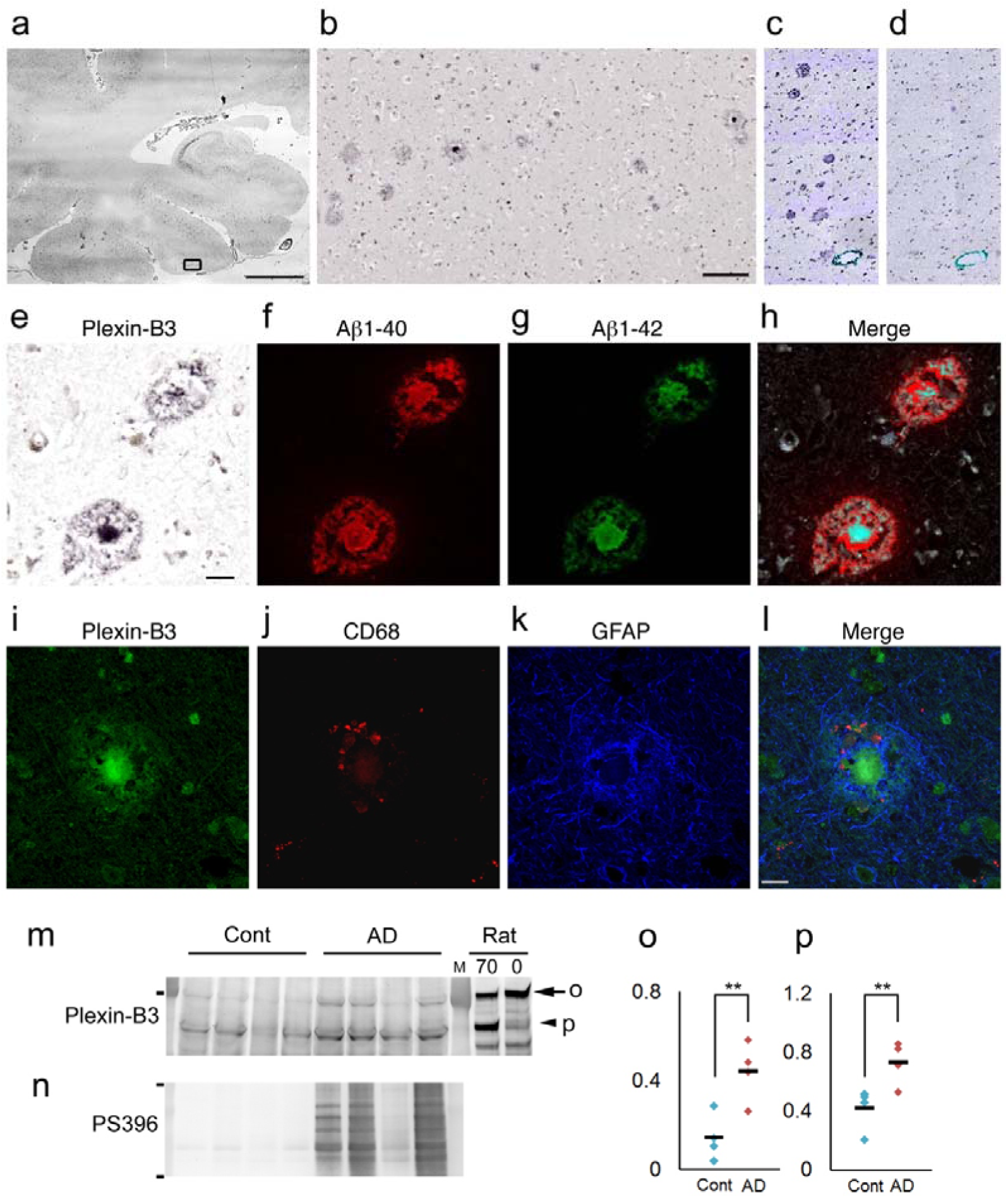
Characterization of plexin-B3^+^ senile plaques (SPs) in AD brains. (a) Cortical distribution of plexin-B3^+^ SPs in an AD brain Scale bar: 5 mm. (b) An enlarged image of the rectangular area in (a) (see also Supplementary Fig. 9). Scale bar: 200 μm. (c & d) Specificity of the anti-plexin-B3 polyclonal antibody against human plexin-B3 (See also Supplementary Fig. 11A & B). (e ∼ h) Spatial relationship between plexin-B3^+^ and Aβ^+^ areas as analyzed by the combination of the phase contrast (e) and confocal images (f ∼ h). Scale bar in d: 20 μm. See also Supplementary Fig. 10. (i ∼ l) Fluorescent immunostaining of plexin-B3, CD68 (a marker for microglia) and GFAP in a SP. Scale bar: 20 μm. (m) Increased plexin-B3 levels in the Sarkosyl-soluble fractions of AD brains. An arrow (200 kDa as marked “o”) and an arrowhead (around 130 kDa as marked “p”) indicate two major bands of plexin-B3. Note that the postmortem interval increased the levels of 130 kDa plexin-B3 band in the rat brains. Rat 0: adult rat brain homogenate with 0 hr postmortem interval; Rat 70: adult rat brain homogenate with 70 hrs postmortem interval. Molecular weight marker: 200 kDa. (n) Accumulation of abnormally hyperphosphorylated tau at Ser396 (PS396) in the Sarkosyl-insoluble fractions of the AD brains. Note that abnormal tau accumulation is clearly associated with increased plexin-B3 level. Molecular weight markers: 87 and 35 kDa. (o & p) Quantification of the intensity of the plexin-B3 bands in panel (m). Adjusted arbitrary units were ploted as Y-axis. \*\**P* < 0.03

WB analyses of Sarkosyl-soluble and -insoluble fractions of the frozen normal control and AD brains (Supplementary Table 3B) further revealed that the expression of plexin-B3 was detected only in the Sarkosyl-soluble fractions (Fig. 6m). The specificity of the antibody was further confirmed by WB (Supplementary Fig. 11B & C). We found that the levels of plexin-B3 increased in AD brains in association with the accumulation of abnormally hyperphosphorylated tau at Ser396 in the Sarkosyl-insoluble fractions (Fig. 6m∼p).

## Discussion

To understand in greater depth the functions of an individual cell type in the adult CNS and in disease pathogenesis, homogeneous primary cultures would be indispensable. In the present study, we first established a novel reproducible method to purify and culture aOPCs from the adult rat brain and discovered a novel uncharacterized aOPC marker plexin-B3. We further demonstrated several lines of evidence suggesting that plexin-B3^+^ aOPCs represent one of as-yet-unidentified Aβ-secreting cells in adult CNS cells and most likely in AD. This unique idea is supported not only by the data from the functional analyses of aOPCs in vitro and in vivo, but also by the data from the human brain studies, especially by the specific plexin-B3 immunoreactivity of SPs in AD, one of the most essential neuropathological hallmarks of AD. As far as we know, none of the AD animal models could recapitulate the cored SP pathology, suggesting that simple secretion of Aβs into the extracellular matrix does not sufficiently induce such pathology. Our findings suggest the possibility that Aβs accumulate as cored SPs due to the morphology of the cells secreting them.

Our findings also raised a simple question running counter to the conventional amyloid hypothesis which contends that neurons comprise the dominant and, in fact, sole Aβ-secreting cells in AD. One possible alternative mechanism may involve a type of demyelination or dysmyelination process in the active myelinated areas of aged gray matter in association with increased secretion of Aβs from aOPCs and the formation of cored SPs, thereby promoting axonal dysfunctions, neurological disabilities, and most likely formation of neurofibrillary tangles in neurons (Supplementary Fig. 12A & B). This idea is fairly consistent with Braak’s neuropathological findings^11^{Braak, 1996 #1377} (Braak & Braak, 1996) and the recent findings in the unbaised single-cell transcriptome analyses of AD brains^12^. If our scenario is true, fine control of cortical myelination in association with proper APP processing in the OL lineage cells in aged brains would be one of the essential requirement of effective AD therapy (Supplementary Fig. 12C), although the exact roles of APP in aOPCs, or its processing in OL differentiation, remain completely unknown.

Cortical plexin-B3^+^ aOPC gliosis induced by CSD (Fig. 5c, d & e) may represent another promising therapeutic target for AD. Very mild CSD stimulation (1 or 2 times during 10 min) (Fig. 5d) was sufficient to induce delayed plexin-B3^+^ aOPC gliosis in the remote ipsilateral cortex, suggesting that plexin-B3 is one of the most sensitive glial markers for brain injuries. Interestingly, an uncompetitive pan-NMDA-R blocker, memantine, which has been approved for the treatment of dementia, has proven to be partially protective against CSD^43^. The preventive use of memantine-type derivatives in MCI or during the early phase of AD seems promising.

This study also sheds light on several signal transduction pathways, including FGF2 and plexin-B3-semaphorine, as potential therapeutic targets for AD. For example, our in vitro data clearly indicate that the loss of FGF2 signaling is amyloidgenic in aOPCs (Fig. 3). This, however, does not simply mean that FGF2 replenishment confers therapeutic benefits for AD, since it also promotes aOPC proliferation, thereby inhibiting OL differentiation (Fig. 1d & f) and likely disturbing normal myelination. Temporal up- and down-regulation of FGF2 may also be involved in the induction of plexin-B3^+^ aOPC gliosis in the CSD models^44, 45^. In AD brains, elevated FGF2 levels have been reported^46, 47^, further highlighting the need for clarifying the exact roles played by FGF2 in the pathogenesis of AD.

In contrast, almost nothing is known about the roles of plexin-B3 in physiological OL differentiation^48^ or in AD pathogenesis. Plexin-B3, which is expressed mainly in postnatal brains^49^, is a high-affinity receptor specific for semaphorines^24, 48^. Mice lacking plexin-B3 display normal CNS morphology and behaviour^49^. Even the fate of plexin-B3^+^ aOPCs in CSD-induced cortical gliosis remains unclear at present. Since plexin-B3 act at least as a temporal aOPC marker expressed during the initiation of OL differentiation, plexin-B3^+^ aOPCs may differentiate into mature OLs for the recovery, remodeling, and remyelination of neuronal circuits after cortical injuries. However, we hypothesize that such mechanisms may be largely destroyed in AD (Supplementary Fig. 12). Further studies are clearly needed to understand the fundamental mechanisms of adult cortical myelination, the roles of plexin-B3^+^ aOPCs in health and disease, the pathophysiology of AD-type de- or dysmyelination, and its therapeutic interventions.

In conclusion, we not only provide both a novel culture method for aOPCs and an aOPC marker plexin-B3, but also demonstrate several lines of evidence suggesting that plexin-B3^+^ aOPCs may be one of the major Aβ-secreting cells in AD. The culture method will be useful for discovering novel functions of aOPCs as well as the regulatory mechanisms underlying adult OL differentiation. Furthermore, these findings will shed light on new AD pharmacotherapies targeting plexin-B3^+^ aOPC differentiation and cortical (oligodendro)gliosis.

## Materials and Methods

All protocols were approved by the Tokyo Metropolitan Institute of Medical Science Animal Care and Use Committee.

### aOPC culture

Rats (female Sprague-Dawley (SD) rats, > 2 months old) were deeply anesthetized with pentobarbital (50 mg/kg). The hippocampus and other brain regions were dissected according to the Paxinos and Watson atlas (http://labs.gaidi.ca/rat-brain-atlas/) (see Supplementary Table 1), finely minced and digested in papain (26.1 U/ml) (Worthington, Lakewood, NJ) in Hibernate A (Brainbits, Springfield, IL) containing 2% B27 (Thermo Fisher Scientific, Waltham, MA) and 0.5 mM glutamine (Thermo Fisher Scientific)(referred to as Hibernate A/B27) for 30 min at 30°C with shaking at 170 rpm. The tissue was gently dissociated in 2 ml warm Hibernate A/B27 via brief mechanical trituration. The suspension settled for 2 min, and the resulting supernatant (2 ml) was collected. This series of dissociations was repeated three times. A total of 6 ml of supernatant was gently overlaid on a step gradient composed of four parts of 1 ml Hibernate A/B27 with 16.4% (density 1.041 g/ml), 11.7% (1.029 g/ml), 9.4% (1.023 g/ml) and 7% (1.017 g/ml) Optiprep^TM^ (1.320 g/ml) (Thermo Fisher Scientific). After centrifugation at 800 x *g* for 15 min at room temperature, the top 6 ml of the supernatant was discarded. The remaining 4 ml fractions consisted mainly of two parts: a large dense band in the middle of the 1.029 g/ml fraction (“sup fraction”, or “S” in Figures) and a discrete pellet at the bottom of the tube (“pellet fraction,” or “P”) (Fig.1a).

The sup fraction was collected, mixed with 4 ml of medium consisting of Neurobasal A supplemented with 2% B27, 0.5 mM glutamine, 100 IU/ml of penicillin and 100 μg/ml of streptomycin (Thermo Fisher Scientific) (Neurobasal A/B27), and centrifuged at 800 x *g* for 15 min. The pellet was then resuspended in 10 ml Neurobasal A/B27, and the suspensions were plated onto 60-mm dishes (Corning, Corning, NY) coated with poly-d-lysine (5 mg/50 ml distilled water for 2 hours, PDL; 135 kDa; Sigma-Aldrich, St. Louis, MO) or onto poly-d-lysine-coated BD BioCoat^TM^ (BD Bioscience, San Jose, CA) chambers. The dishes (or chambers) were incubated at 37°C with 5% CO_2_ for 30-60 min and then gently washed to remove unattached cells and debris. New Neurobasal A/B27 containing 20 ng/ml FGF2 (Peprotech, Rocky Hill, NJ) was added to the dishes (4 ml) or chambers (400 μl). Half of the medium was changed to new medium with a double amount of fresh FGF2 added every 2-3 days. The cells were passaged at 70-90% confluence.

### Immunocytochemistry

The cells were fixed with 4% paraformaldehyde in PBS for 15 min. After permeabilization with 0.5% Triton X-100 in PBS for 5 min, the cells were blocked in 3% bovine serum albumin (BSA)(Sigma-Aldrich) in TBS containing 0.05% Tween 20 (0.05% TBS-T) for 20 min and incubated with the primary antibodies (Supplementary Table 2) overnight at 4°C and followed by Alexa 488-, Alexa 594-, or Alexa 630-labeled secondary antibodies, TO-PRO-3 (Thermo Fisher Scientific), or Hoechst 33258 in the blocking buffer for 60 min.

### Western blot

The samples were lysed with modified-RIPA buffer (50 mM Tris-HCl pH 8.0, 150 mM NaCl, 1% NP-40, 0.25% sodium deoxycholate, 1 mM EGTA pH 7.4) containing a protease inhibitor cocktail (Sigma-Aldrich). After sonication, the lysates were centrifuged at 15,000 x *g* for 10 min at 4°C. The resulting supernatants were collected, and the protein concentrations were determined using the BCA method (Thermo Fisher Scientific). The samples were separated on a 5–20% gradient SDS-PAGE (Wako, Osaka, Japan) and transferred. The blots were blocked with 1% BSA (Sigma-Aldrich) in 0.05% TBS-T (blocking buffer) for 1 hour, and the membranes were incubated with primary antibodies (Supplementary Table 2) in blocking buffer overnight, followed by secondary antibodies conjugated with HRP or alkaline phosphatase (Promega, Madison, WI, 1:1,000). Proteins were visualized with a LAS-3000 mini (GE Healthcare, Hino, Japan) using enhanced chemiluminescence (Chemi-Lumi One) (Nakalai, Kyoto, Japan), and then visualized with a BCIP-NBT Solution Kit (Nakalai).

Quantitative measurements were performed using Multi Gauge V2.3 software (GE Healthcare). In general, measured protein signal was subtracted from the backgraound signal. For the WB analysis of human brains, the final signal was further adjusted by the signal of the each defined lane of the Ponceau S staining (Nakalai).

### RNA extraction

Cultured aOPCs were washed with cold PBS and frozen at −80°C until shipping. Total RNA was purified using an RNeasy Mini Kit (QIAGEN, Limburg, Netherlands) in accordance with the manufacturer’s instructions. RNA quality was accessed by Bioanalyzer analysis.

### Microarray

Samples were shipped to Agilent Array Services (Hokkaido System Science, Sapporo, Japan). RNA was amplified into cRNA and labeled according to the Agilent One-Color Microarray-Based Gene Expression Analysis Protocol (Agilent Technologies, Santa Clara, CA). The samples were hybridized to Rat GE 4×44K v3 array slides, and the arrays were then scanned using an Agilent Microarray Scanner (Agilent Technologies). The scanned images were analyzed using the standard procedures described in the Agilent Feature Extraction software 9.5.3.1 (Agilent Technologies).

To compare rat and mouse genes, rat orthologs were checked manually through PubMed (http://www.ncbi.nlm.nih.gov/pubmed) or the Rat Genome Database (http://rgd.mcw.edu/). We excluded genes from the lists if no rat ortholog of the gene was found or if it was not listed in the Agilent Rat GE 4×44K v3 array. When more than one expression data was obtained for a gene, the largest expression data was used. Complete lists of the cell-type-specific genes appear in Supplementary Table 4 – 6, and the top genes appear in Fig. 1f & Supplementary Fig. 2.

### RNA-sequencing

RNA extraction and RNA-sequencing (RNA-seq) were performed in TaKaRa RNA-seq Services (TaKaRa, Kusatsu, Japan), including library preparation, fragmentation and PCR enrichment of target RNA. Samples with an RNA integrity number greater than 8 were used for library construction. Sequencing libraries were prepared using a TruSeq RNA Sample Prep Kit (Illumina) according to manufacturer’s instructions and then sequenced by the HiSeq 2500 platform (Illumina) to obtain 100 bp paired-end reads.

One hundred bp paired-end reads were aligned to the rat reference genome (University of California, Santa Cruz (UCSC) Genome Browser Jul.2014 (RGSC 6.0/rn6)) using TopHat tool (version 2.0.14), which incorporates the short read aligner Bowtie (version 2.2.5). We assessed genes whose estimated fragments per kilobase of transcript per million mapped reads (FPKM) values had transcripts with an expression of > 0 FPKM as “TRUE” and the other FPKM as “FALSE”. Complete lists of the cell-type-specific genes appear in Supplementary Table 7 – 9 and the top genes appear in Supplementary Fig. 3 & 4A – 4C.

### Immunoprecipitation

A Pierce Classic IP Kit (Thermo Fisher Scientific) was used for the Aβ immunoprecipitation. Briefly, conditioned medium (3.6 ml) was adjusted to a 1:1,000 protease inhibitor mixture (Sigma-Aldrich) and then incubated with combinations of antibodies (Aβ1-40 (2 μg) or Aβ1-42 (2 μg) + APP (22C11, 5 μg) and with 20 μl Protein A/G resin at 4°C overnight.

### ELISA

A two-site sandwich ELISA^50^ was also used for the measurement of Aβ levels. Cultures of embryonic rat hippocampal cells were prepared as previously described^51^.

### Image analysis

Fluorescent images were observed using high-resolution confocal microscopies (Zeiss, Oberkochen, Germany). In some cases, the entire area of the immunostained section was digitalized with a virtual slide system (VS120) (Olympus, Tokyo, Japan). Plotting was performed manually on transparent layers, overlaid onto the original virtual slide images (x 10 objective). For cell counting, defined areas (e.g., Supplementary Fig. 8) were captured by a microscope equipped with AxioCam MRc 5 (x 20 objective) (Zeiss).

For the quantitative immunofluoresent histogram, cultured cells were imaged and analyzed using the Zen 2011 imaging software (Zeiss). For definition of the region of interest (ROI), the whole cell territory was first outlined in light blue line under the phase contrast image and then lacked areas outlined by yellow lines were subtracted (Fig. 3f). The signal intensity of the ROI was measured and adjusted by the area (= Mean Intensity).

### Animal studies

SD rats (Charles River Laboratories, Yokohama, Japan) were deeply anesthetized with pentobarbital (50 mg/kg) and placed in a stereotactic frame.

For stab wound injuries, 4 male animals (3 months) were underwent a stab wound in the right cortex (Bregma AP −3.6 mm, ML 2.4 mm). Rats were killed at 2 (n = 2) and 7 (n = 4) days after the lesion.

For KCl injuries, animals were assigned to the following groups: sham-operated controls (n = 2), 2 ∼ 3-days (2D, n = 3), 6 ∼ 7-days (7D, n = 4), or 17-days (n = 3) after 3 M KCl, and 3-days (n = 1) and 7-days (n = 1) after 3 M NaCl treatment. A right parietal trepanation (AP −5.7 mm, L 2.7 mm) was carried out and the dura was incised for the topical application of a cotton ball soaked with 3 M KCl or NaCl. After 10 min, the application site was rinsed with saline and the animals were allowed to survive.

To detect proliferating cells in vivo, 10 mg/ml BrdU was administered intraperitoneally (50 mg/Kg) twice/day for 5 consecutive days.

For immunohistochemistry, the animals were anesthetized and transcardially perfused with PBS followed by 4% paraformaldehyde in PBS. Brains were removed, post-fixed for 1 day at 4°C, and then cryoprotected overnight in 20% sucrose in PBS. Coronal sections were cut with a freezing microtome at 40 µm. Sections were generally pretreated with DAKO Target Retrieval Solution (pH 9.0) (Agilent Technologies) and then incubated overnight at 4°C in primary antibodies (Supplementary Table 2) in PBS containing 5% goat serum (or BSA) and 0.4% Triton X-100, followed by 2 hours at room temperature in corresponding secondary antibodies with or without TO-PRO-3 or Hoechst 33258. For observation under a light field, the sections were incubated with streptavidin biotinylated horseradish peroxidase complex (ABC Elite) (Vector, Burlingame, CA). Color development was performed with 3,3’-diaminobenzidine (Sigma-Aldrich) in the presence of imidazole and nickel ammonium chloride. Extracellular DC field potential recordings were made using 0.3-mm-diameter Ag/AgCl microelectrodes (TN217-001) (Unique Medical, Tokyo, Japan). The rinsed recording electrode was positioned on the cortical surface (0.3mm anterior and 2.7mm lateral to bregma) after removal of the dura mater. An Ag/AgCl reference electrode (TN217-002) (Unique Medical) was placed on the cerebellum surface. Electrodes were connected to a differential headstage and amplified using a differential Extracellular Amplifier (gain of 100, ER-1) (Cygnus Technology, Delaware, PA). The high cut filter was set at 300 Hz. DC potentials were digitized at a sampling frequency of 1kHz with a CED Power1401 data-acquisition system and Spike2 software (Cambridge Electronic Device, Cambridge, UK).

### Human studies

The autopsied human brains (Supplementary Table 3A) were fixed in 10% formalin and embedded in paraffin wax. Five to 20-μm-thick sections from the hippocampus were obtained. For plexin-B3 immunohistochemistry, serial pretreatment with heat (110°C in 0.01 M citrate buffer for 10 min) and trypsin (0.05 % for 10 min) was performed. Samples were then incubated in primary antibodies diluted in PBS containing 5% BSA and 0.03% Triton X-100 at 4°C for overnight, and biotinylated secondary antibodies (1:1000) (Vector) for 2 hours. After subsequent incubation with streptavidin biotinylated horseradish peroxidase complex (Vector), color development was performed. For 4G8, Aβ1-42, and plexin-B3 triple immuno(fluoro)labeling, the sections were first immunostained for plexin-B3 as described above. After the digitalized images were obtained under a light field, the sections were further treated with formic acid (>99 % for 1 min) for Aβ double immunofluorolabelling (4G8 and Aβ1-42). To investigate the specificity of the sheep polyclonal antibody for plexin-B3 (R&D systems, Minneapolis, MN), the antibody was pretreated with or without 5x recombinant human plexin-B3 (His45-Gln1255 (Gln1156Asp) with a C-terminal 6-His tag) (R&D systems) overnight.

For WB analysis of human brains (Supplementary Table 3B), biochemical fractionation from frozen postmortem brain homogenates was performed as previously described^52^. Briefly, unfixed frozen human brain blocks (Brodmann area 6) were homogenized in 9 volumes of extraction buffer A68 (10 mM Tris–HCl, pH 7.5, 1 mM EGTA, 10% sucrose, 0.8 M NaCl) containing a protease inhibitor cocktail (Calbiochem, San Diego, CA). After adding 20% (w/v of water) Sarkosyl to concentrations of 2% w/v, the homogenates were left for 30 min at 37 °C, followed by a 10min spin at 15,000rpm. The resulting supernatants were further centrifuged at 50,000rpm for 20min. The final supernatants retained as Sarkosyl-soluble fractions and the pellets were solubilized in PBS as Sarkosyl-insoluble fractions. To examine the effects of postmortem interval on the plexin-B3 expression, frozen rat brains dissected from the rats with 0 hr and with 70 hrs (at 4 °C) postmortem intervals were fractionated and analyized as described above.

### Data analysis

The data are expressed as the mean ± SD unless otherwise indicated. Statistical comparisons were performed using paired Student’s *t*-tests.

## ACKNOWLEDGEMENTS

We thank T. Tomita for technical advice. We thank T. Shinozaki, H. Kondo, and A. Nakamura for their technical assistance. This work was supported by JSPS KAKENHI Grant Numbers 25461794 for N.N.-K. and 15K15438 for Y.T.

## AUTHOR CONTRIBUTIONS

Y.T. designed the study; N.N.-K., X.-J. Y., T.I., A.T. and Y.T. established the culture model, collected and processed the data; Y.M. and N.O. conducted animal surgeries and N.N.-K. and Y.T. analyzed the animal data; S.T. and T.U. collected human samples; S.T., T.U., N.N.-K., M.H. and Y.T. analyzed the human data; K.S. T.K. and Y.T. analyzed the microarray and RNA-seq data; Y.T. wrote the paper; All authors discussed the results and commented on the manuscript.

## CONFLICT OF INTERST

The authors declare no competing financial interests.

