## Supplementary Information for "Essential roles of plexin-B3^+^ oligodendrocyte precursor cells in the pathogenesis of Alzheimer’s disease"

Naomi Nihonmatsu-Kikuchi et al.

**Contents:**

**Supplementary Fig. 1 - 12**

**Supplementary Table 1 - 3**

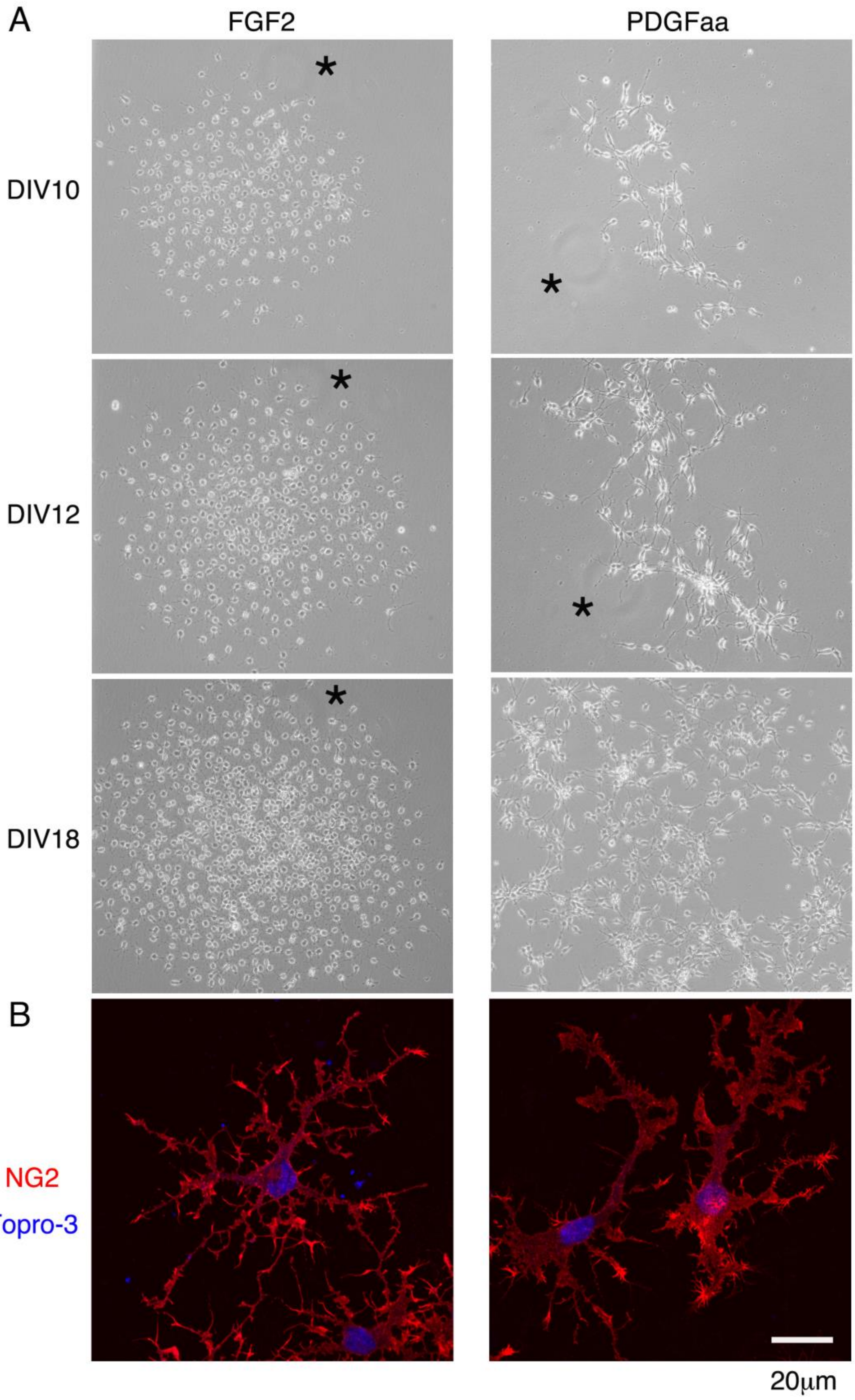
A
FGF2
PDGFaa
DIV10
DIV12
DIV18
*
B
NG2
Topro-3
20μm

**Supplementary Fig. 1. Effects FGF2 or PDGF-aa on NG2+ aOPCs in the primary cultures.**

(**A**) Phase contrast images of primary cultures in FGF2 (20 ng/ml) or PDGF-aa (20 ng/ml): The phase contrast images at 10, 12 and 18 days in vitro (DIV) show the typical colony formations in FGF2 or PDGF-aa. While cells in FGF2 migrated slowly, those in PDGF-aa migrated relatively faster. While cells in FGF2 were of high density, those in PDGF-aa were relatively scattered. Note that, in the passaged cultures, PDGFaa no longer renders NG2+ aOPCs proliferative (see **Fig. 1d**). Asterisks indicate position markers. Scale bar: 300 μm. (**B**) Morphology of NG2+ cells in FGF2 (20 ng/ml, Left) or PDGF-aa (20 ng/ml, Right): The cells were immunostained with anti-NG2 antibody (red) and TO-PRO-3 (blue). The typical morphologies of NG2+ OPCs in FGF2 were multipolar, while those in PDGF-aa were generally bipolar. Scale bar: 20 μm.

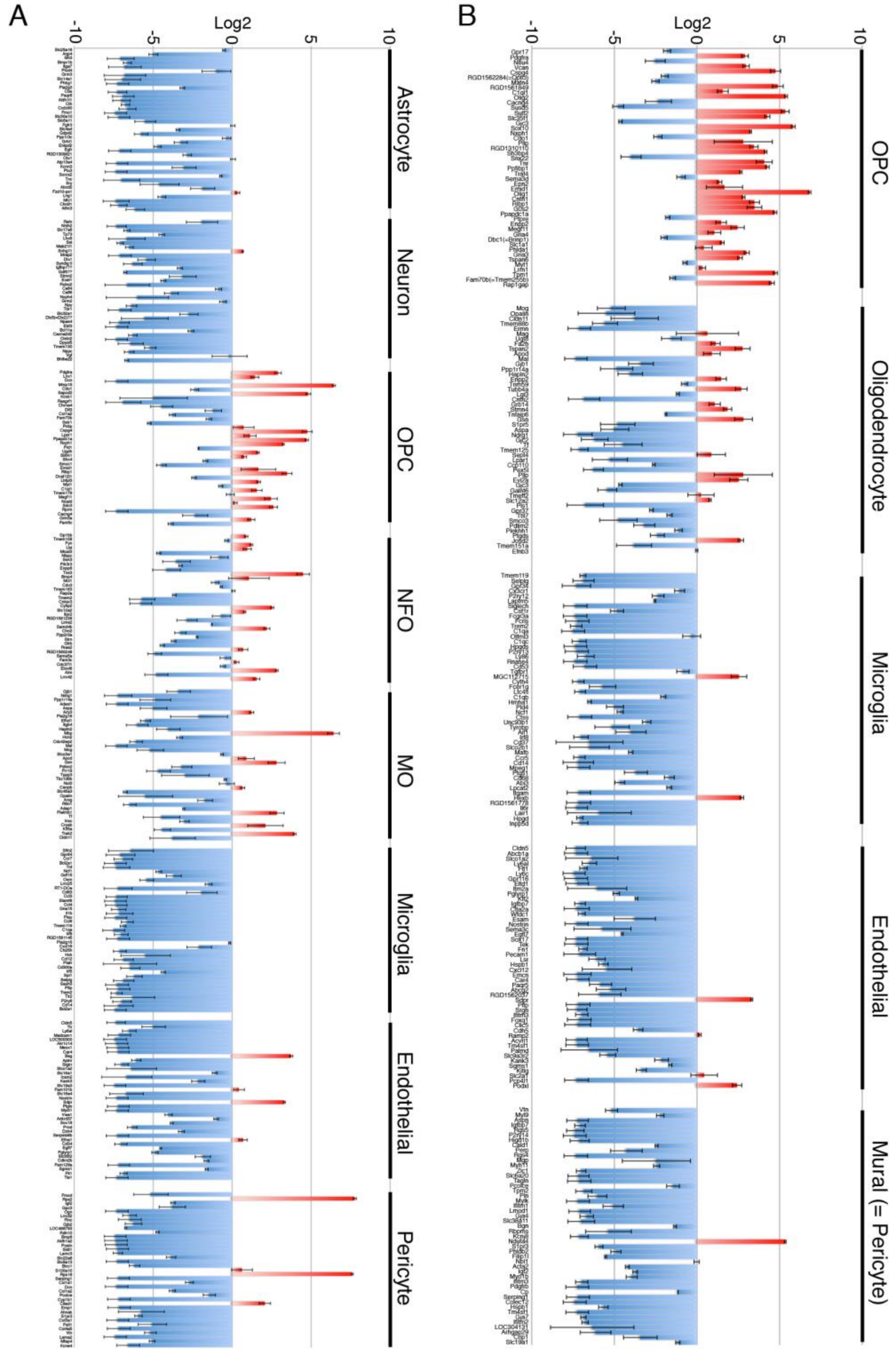


**Supplementary Fig. 2. Microarray profiles of cultured aOPCs.**

(**A**) Comparison with the top 40 cell type-specific genes listed in Zhang et al. (2014): The top 40 genes of astrocyte, neuron, OPC, NFO (newly formed oligodendrocyte), MO (myelinating oligodendrocyte), microglia, endothelial cell, and pericyte from adult mouse cortex by Zhang et al. (2014) were compared with the microarray data. Note that cultured aOPCs generally express oligodendrocyte lineage (OPC, NFO, and MO) specific-genes, especially those specific to OPC. Although the reason is unclear, two pericyte specific genes, ribosomal protein S2 (Rps2) and Rsp18, were highly enriched in cultured aOPCs. For the complete list of the genes, see also Supplementary Table 5. (**B**) Comparison with the top 50 genes listed in Wu et al. (2017): Top 50 genes of OPC, oligodendrocyte, microglia, endothelial cell, and mural cell (pericyte) obtained from single-cell RNA-seq of 8-10 month-old mouse amygdala by Wu et al. (2017) were compared with the microarray data. See also Supplementary Table 6.

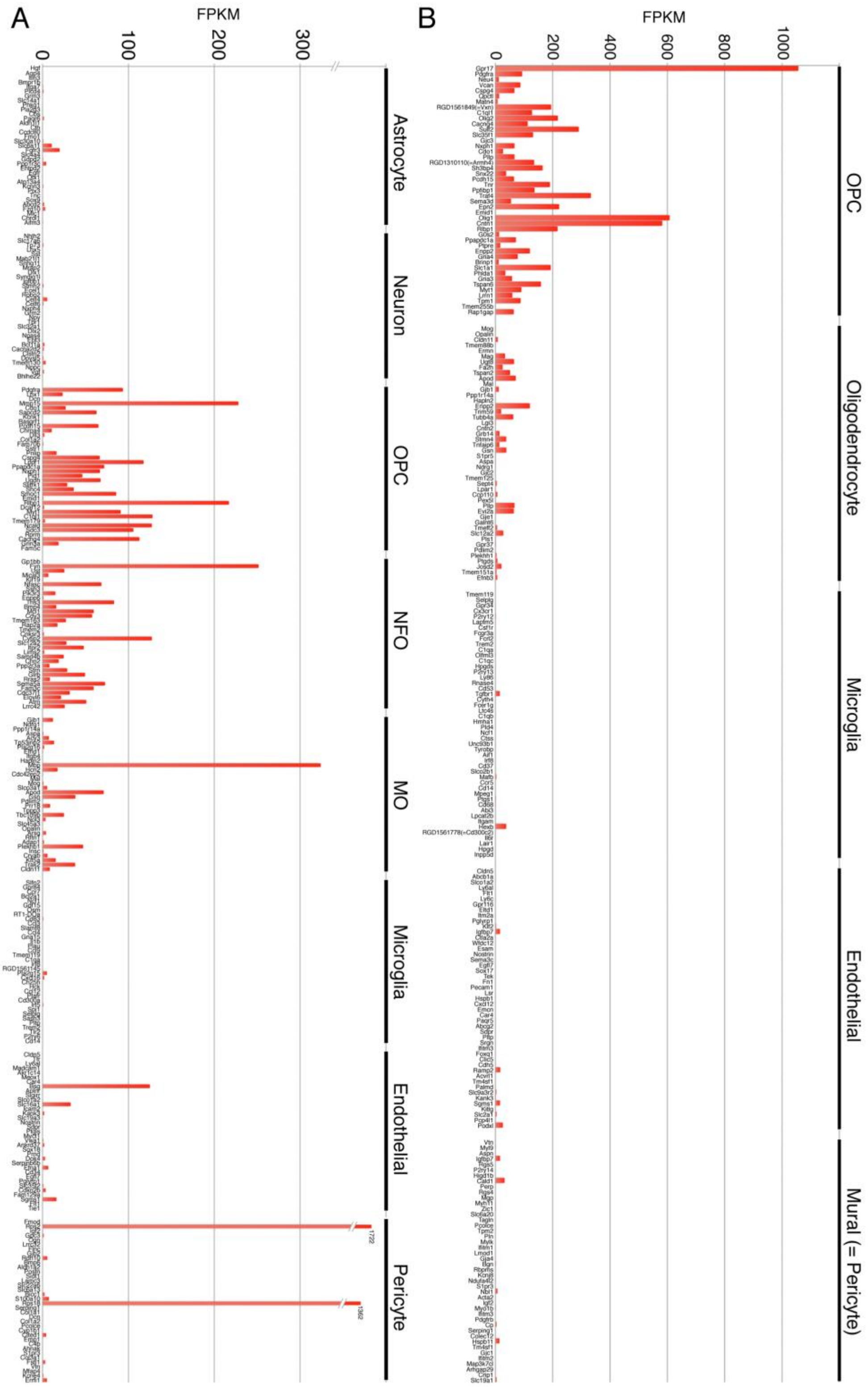


**Supplementary Fig. 3. RNA-seq profiles of cultured aOPCs.**

(**A**) Comparison with the top 40 cell type-specific genes listed in Zhang et al. (2014): The top 40 genes of astrocyte, neuron, OPC, NFO (newly formed oligodendrocyte), MO (myelinating oligodendrocyte), microglia, endothelial cell, and pericyte from the adult mouse cortex by Zhang et al. (2014) were compared with the RNA-seq data obtained from cultured aOPCs. Note that cultured aOPCs express OPC specific-genes. As observed in the microarray analysis, two pericyte specific genes, Rps2 and Rsp18, were highly enriched in cultured aOPCs. See also Supplementary Table 8. (**B**) Comparison with the top 50 genes listed in Wu et al. (2017): Top 50 genes of OPC, oligodendrocyte, microglia, endothelial cell, and mural cell (pericyte) obtained from single-cell RNA-seq of 8-10 month-old mouse amygdala by Wu et al. (2017) were compared with the RNA-seq data obtained from cultured aOPCs. Note that cultured aOPCs express OPC specific-genes. See also Supplementary Table 9.

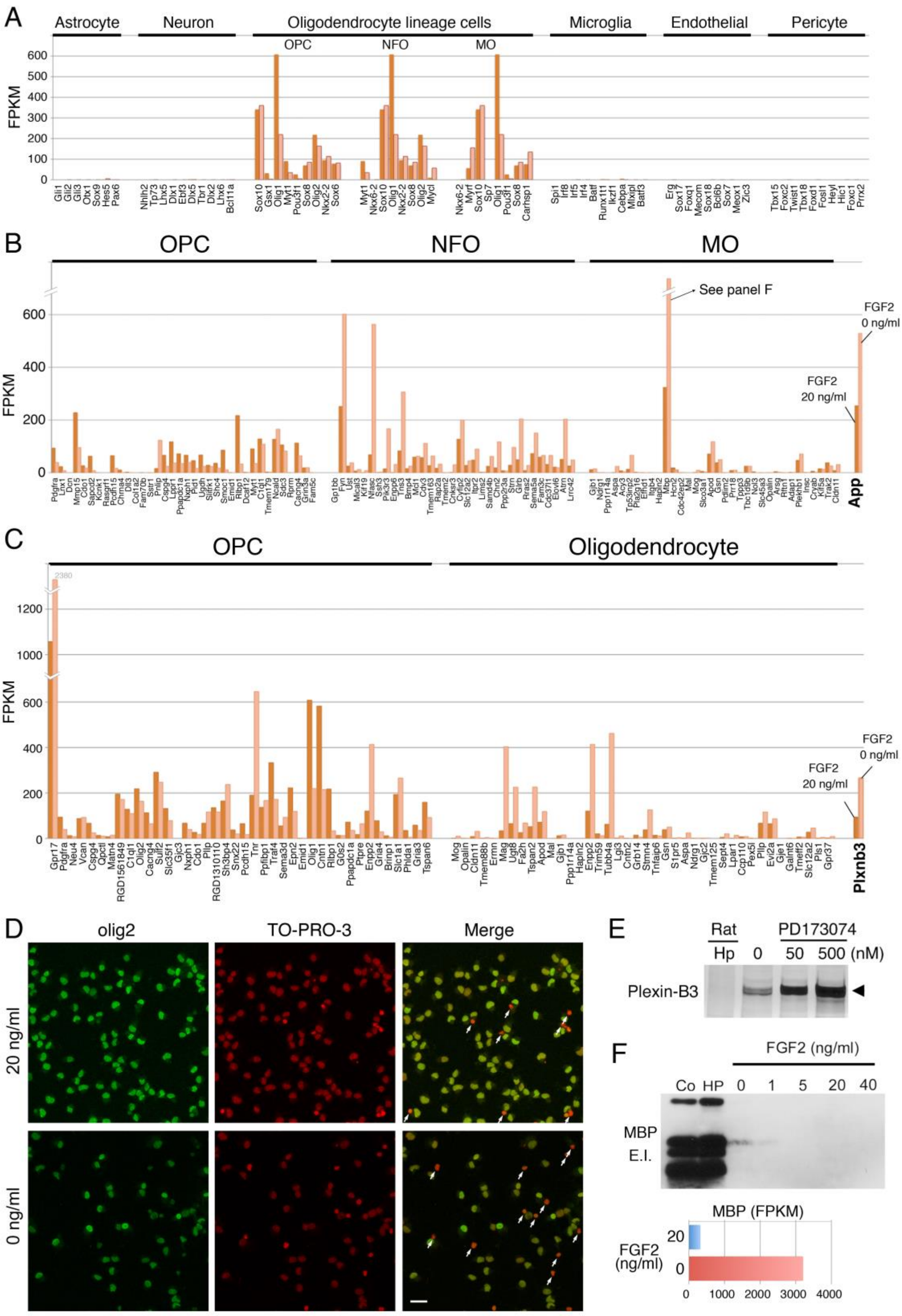

A
Astrocyte
Neuron
Oligodendrocyte lineage cells
OPC
NFO
MO
Microglia
Endothelial
Pericyte
FPKM
B
OPC
NFO
MO
See panel F
FGF2 0 ng/ml
FGF2 20 ng/ml
FPKM
App
C
OPC
Oligodendrocyte
FPKM
FGF2 20 ng/ml
FGF2 0 ng/ml
Plxnb3
D
olig2
TO-PRO-3
Merge
20 ng/ml
0 ng/ml
E
Rat
Hp
PD173074
0
50
500 (nM)
Plexin-B3
F
FGF2 (ng/ml)
Co
HP
0
1
5
20
40
MBP
E.I.
MBP (FPKM)
FGF2 (ng/ml)
20
0
0
1000
2000
3000
4000

**Supplementary Fig. 4. Effects of FGF2 withdrawal on cultured aOPCs.**

(**A**) Effects of FGF2 withdrawal on the transcription factor gene expression profiles of cultured aOPCs: The top 10 transcription factor genes of astrocyte, neuron, OPC, NFO (newly formed oligodendrocyte), MO (myelinating oligodendrocyte), microglia, endothelial cell, and pericyte by Zhang et al. (2014) were compared with the RNA-seq data obtained from aOPCs cultured with or without 20 ng/ml FGF2. Note that the 5-day FGF2 withdrawal did not change the transcription factor genes. (**B**) Effect of 5-day FGF2 withdrawal on the RNA-seq profiles of cultured aOPCs: The top 40 genes (OPC, NFO and MO) in Zhang et al. (2014) were compared with the RNA-seq data of cultured aOPCs with or without 20 ng/ml FGF2. Note that 5-day FGF2 withdrawal increased several NFO-specific genes and one MO-specific gene (MBP) in cultured aOPCs (See also panel F). For the complete list of the genes, see Supplemenary Table 8. In the right-most bars, the change in App gene expression is shown. (**C**) Effect of 5-day FGF2 withdrawal on the RNA-seq profiles of cultured aOPCs: The top 40 genes (OPC and oligodendrocyte) in Wu et al. (2017) were compared with the RNA-seq data of cultured aOPCs with or without 20 ng/ml FGF2. Note that 5-day FGF2 withdrawal increased or decreased several OPC- and OL-specific genes in cultured aOPCs. See also Supplemenary Table 9. The change in Plxnb3 gene expression is shown in the right-most bars. (**D**) Effect of FGF2 withdrawal on the proportions of olig2+ cells: Immunocytochemistry of aOPCs cultured with or without 20 ng/ml FGF2 for 5 days is shown. White arrows indicate dead cells that were strongly TO-PRO-3 positive. More than 99% of living cells were olig2+ under both conditions (see also **Fig. 2c**). (**E**) Effects of the FGF inhibitor, PD173074, on the levels of plexin-B3 protein: To analyze the effects of FGF2, the cells were cultured in various concentrations of FGF2 (0, 1, 5, 20, or 40 ng/ml) for 5 days or FGF2 inhibitor PD173074 (0, 50, or 500 nM in the presence of 20 ng/ml FGF2) (Merck, Darmstadt, Germany) for 24 hrs. PD173074 dose-dependently increased plexin-B3 protein levels in cultured aOPCs. Co: adult rat cortex, HP: adult rat hippocampus. (**F**) Effect of FGF2 withdrawal on MBP protein and RNA expression: FGF2 withdrawal for 5 days was not enough to increase MBP protein expression although RNA levels increased about 10-fold. See also panel B. Co: rat cortex, HP: rat hippocampus.

A B

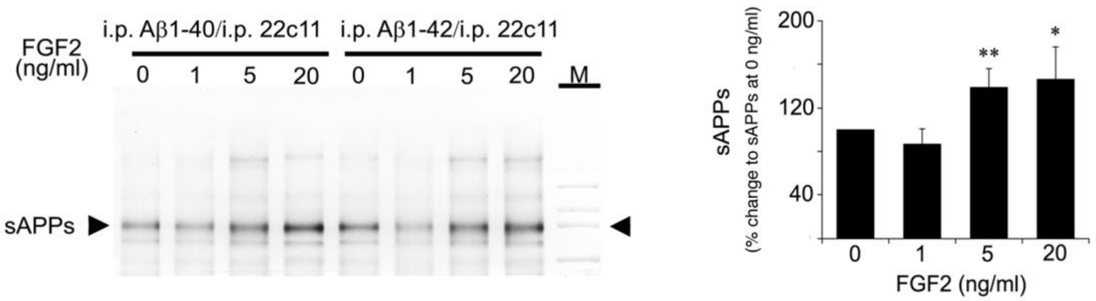


**Supplementary Fig. 5. Effects of FGF2 on APP metabolism in cultured aOPCs.**
(**A**) WB analysis of sAPP in the conditioned media of aOPCs: In parallel with Aβx-40 or Aβx-42 immunoprecipitation (i.p.) (**Fig. 3c**), sAPP, which was liberated from APP by α- or β-secretases, was immunoprecipitated with anti-APP (22c11) antibody from the conditioned media of aOPCs cultured in 0, 1, 5 or 20 ng/ml FGF2 for 24 hrs. The resulting immunoprecipitants were analyzed by WB with 22c11 antibody to quantify the amounts of sAPP. The arrowheads indicate the predicted sAPP band positions (at around 85 kDa). M: molecular markers. (**B**) Quantification of sAPP in the conditioned media: ***P* <0.5x10-6 or **P* <0.03 (in comparison with 0 ng/ml).

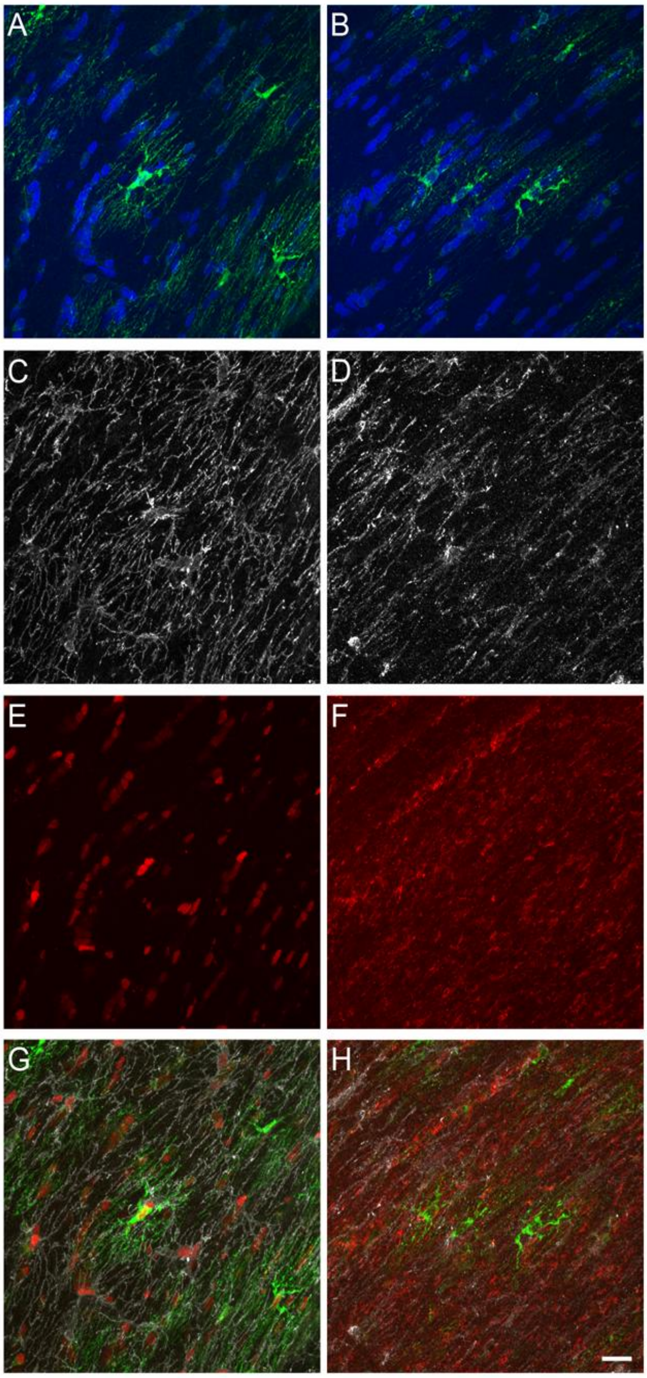


**Supplementary Fig. 6. Plexin-B3+ aOPCs in the rat corpus callosum.**

Brain sections from normal adult rats were immunostained with anti-plexin-B3 (**A** & **B**), anti-NG2 (**C** & **D**), and anti-olig2 (**E**) or anti-MBP (**F**) antibodies. Merged images are shown in **G** & **H**. Nuclear stainings (Blue, Hoechst 33258) are also shown in **A** & **B**. In the corpus callosum, plexin-B3+ aOPCs were negative for MBP and NG2, but positive for olig2. Scale bar: 20 μm.

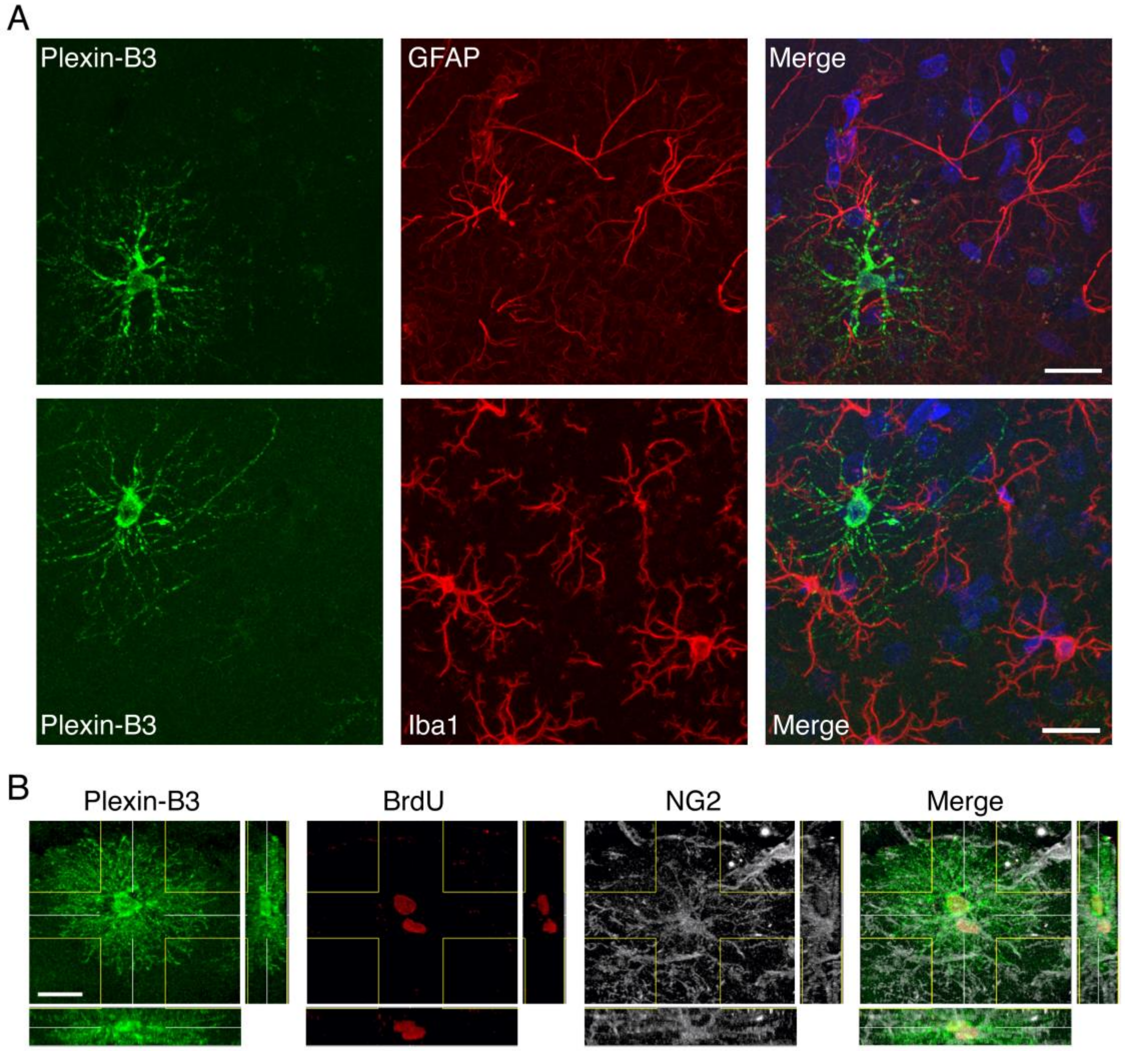


**Supplementary Fig. 7. Plexin-B3+ aOPCs in the rat cortex.**

(**A**) Plexin-B3+ aOPCs were negative for GFAP and Iba1. Brain sections from normal adult rats were immunostained with anti-plexin-B3 and anti-GFAP or anti-Iba1 antibodies. Nuclear stainings (blue, Hoechst 33258) are shown in the "Merge" panels. Scale bar: 20 µm. (**B**) A pair of plexin-B3+/BrdU+ and NG2+/BrdU+ cells most likely formed by asymmetrical aOPC division. Brain sections from normal adult rats treated with BrdU were immunostained with anti-plexin-B3, anti-BrdU, and anti-NG2 antibodies. Merged image (Merge) is shown in the right-most panel. Scale bar: 20 µm.

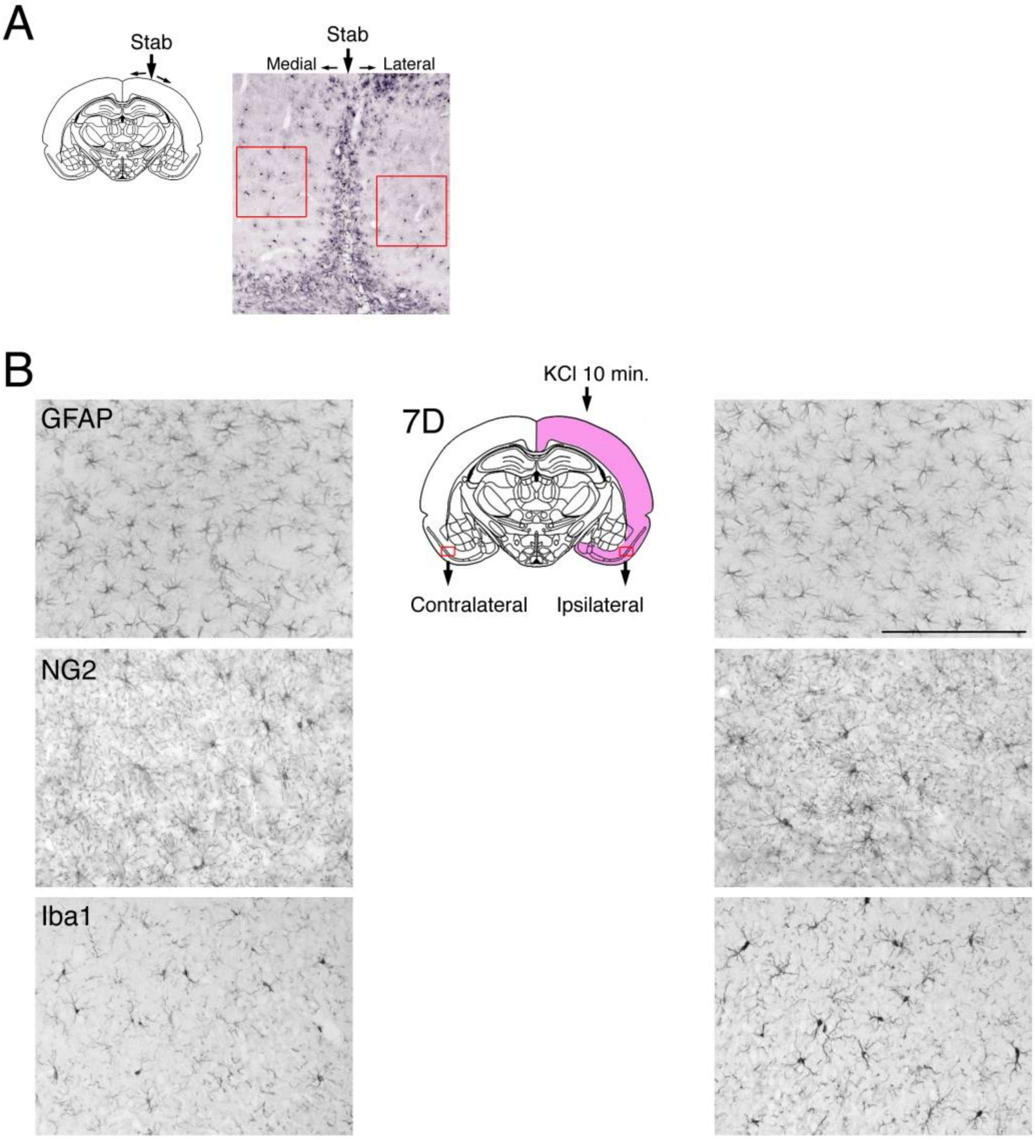


**Supplementary Fig. 8. Areas for quantitative cell counting in the brain injury models.**

(**A**) For the stab wound model, two 200 μm squares were defined in the cortex, about 30 ~ 50 μm apart medially and laterally from the stab lesion. (**B**) For the KCl injury model, two 440 μm x 330 μm rectangles were defined in the ipsilateral and contralateral remote cortex as shown in the schematic figure. At indicated days (0, 2, or 7D) from the injuries, animals were fixed and brain sections were immunostained with antibodies for plexin-B3, GFAP, NG2, or Iba1. After the digital images of the corresponding areas were obtained, cell numbers were counted manually in the defined areas. Scale bar: 200 μm.

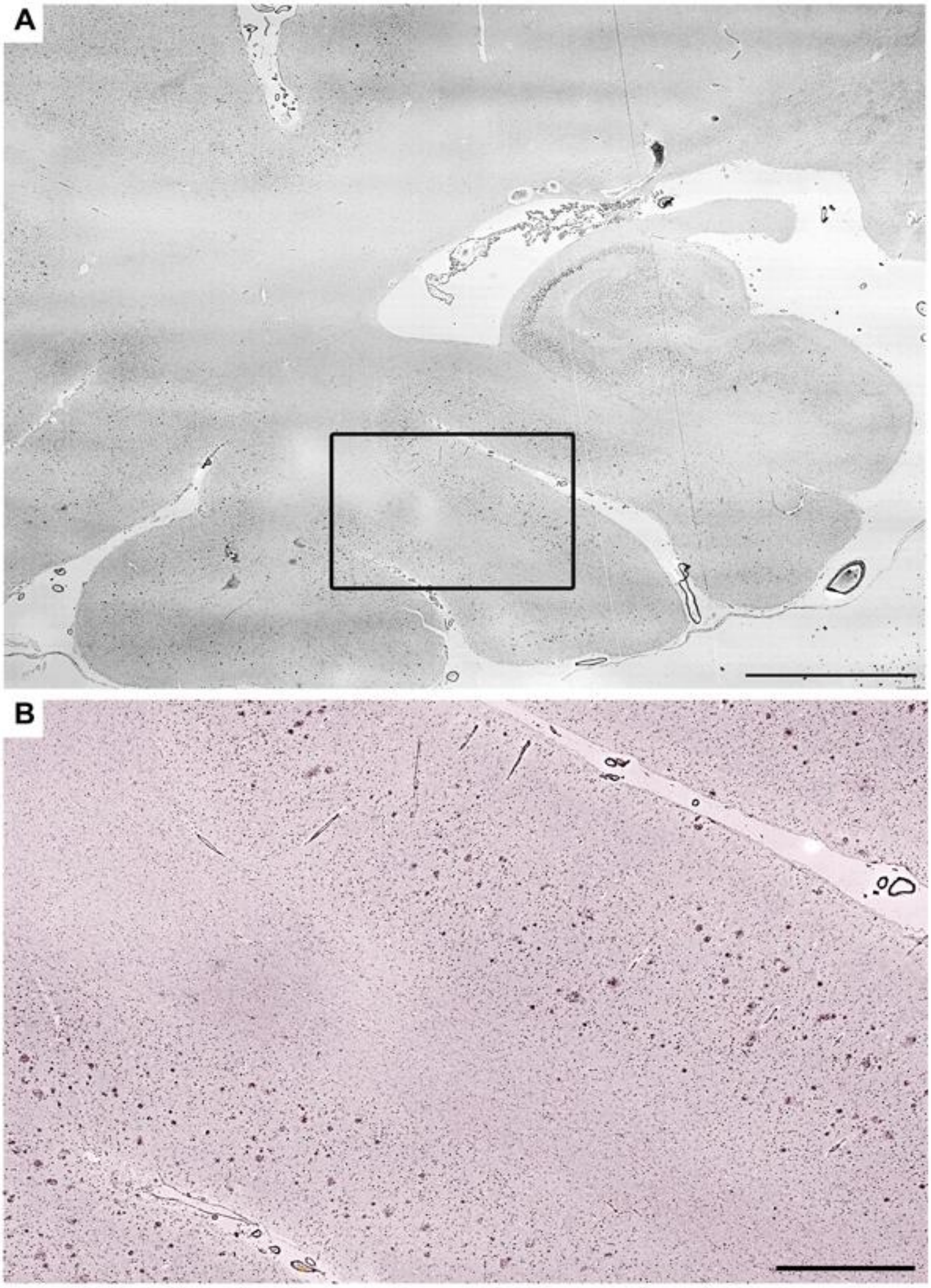


**Supplementary Fig. 9. Cortical distribution of plexin-B3+ senile plaques in the Alzheimer's disease brain**

Immunohistochemistry of an Alzheimer's disease brain with anti-plexin-B3 polyclonal antibody. Plexin-B3 antibody stained mainly cortical structures (**A** & **B**), except for dot-like structures distributed throughout the entire brain (**B**). Scale bar in **A**, 5 mm: Scale bar in **B**: 1 mm.

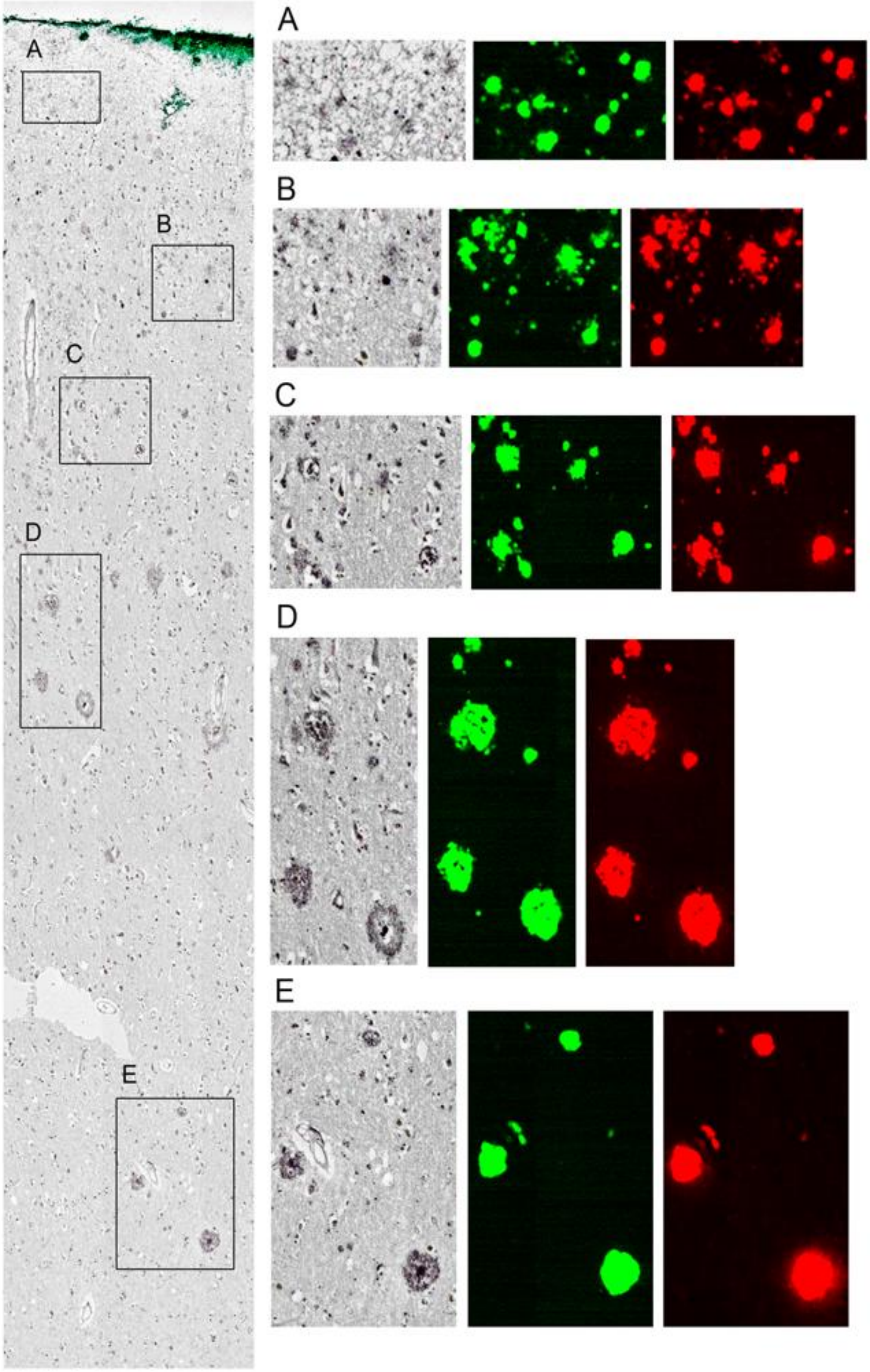


**Supplementary Fig. 10. Spatial relationship between plexin-B3+ and Aβ+ areas.** Immunostaining of an Alzheimer's disease brain with anti-plexin-B3 polyclonal antibody and antibodies for total Aβ (4G8, green) and Aβ1-42 (red). Images were digitalized with a virtual slide system (VS120, Olympus, Tokyo, Japan). Almost all the SPs were co-immunolabelled with anti-plexin-B3 antibody. Scale bar in the left-most panel, 200 μm: Scale bar in **A**: 20 μm: Scale bars in **B**, **C**, **D**, and **E**: 50 μm.

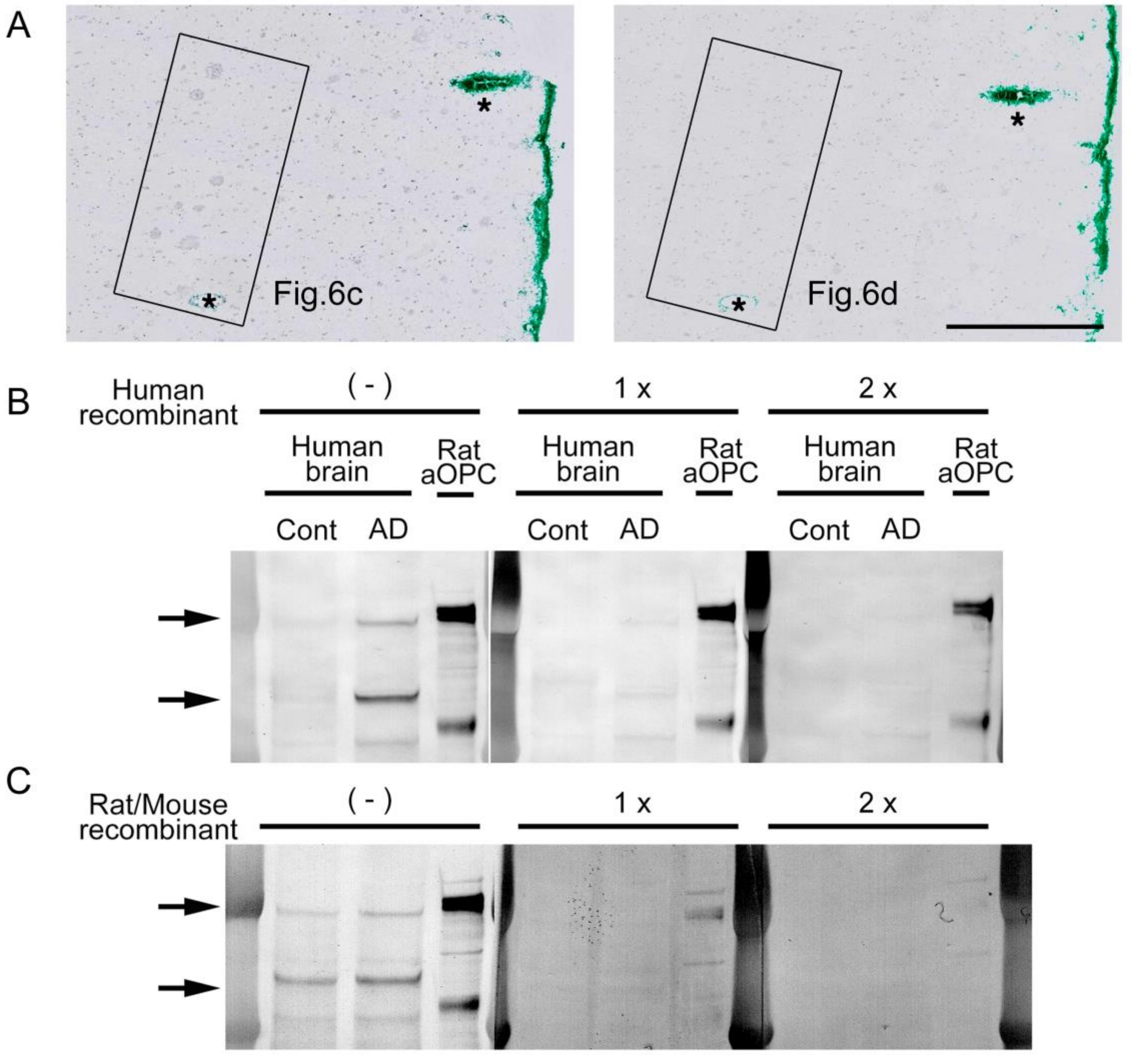


D

| | Human | Rat |
|---|---|---|
| Human recombinant | ✓ | × |
| Rat/Mouse recombinant | ✓ | ✓ |

**Supplementary Fig. 11. Specificity of the anti-plexin-B3 polyclonal antibody.**

(**A**) Original images of Figure 6c and d (as indicated Fig. 6c and d in the panels respectively). Serial sections from an Alzheimer's disease brain were immunostained with anti-plexin-B3 polyclonal antibody (R&D) pretreated without (left) or with (right) 5x recombinant human plexin-B3 (His45 – Gln1255 (Glu1156Asp) with a C-terminal 6-His tag) (R&D) overnight and visualized. Note that pre-absorption successfully eliminated the plaque, but not the dot-like stainings, suggesting that the latter is most likely non-specific. Asterisks indicate blood vessels colored by a green marker before sectioning. Scale bar: 100 μm. (**B**) WB analysis for the specificity of the anti-plexin-B3 polyclonal antibody (R&D) against human plexin-B3. The antibody was pretreated without (-) or with 1x or 2x recombinant human plexin-B3 (His45 – Gln1255 (Glu1156Asp) with a C-terminal 6-His tag) (R&D) overnight and used for the WB analysis. Note that two major bands of human plexin-B3 in the Sarkosyl-soluble fractions, but not rat plexin-B3 expressed in the cultured aOPCs, were successfully eliminated by the pre-absorption. (**C**) WB analysis for the specificity of the anti-plexin-B3 polyclonal antibody (R&D) against rat plexin-B3. The antibody was pretreated without (-) or with 1x or 2x recombinant Rat/Mouse plexin-B3 (His25-Gln1235 with a C-terminal 6-His tag) (R&D) overnight and used for the WB analysis. Note that Both human and rat plexin-B3 was successfully eliminated by the pre-absorption. (**D**) Summary of the pre-absorption studies.

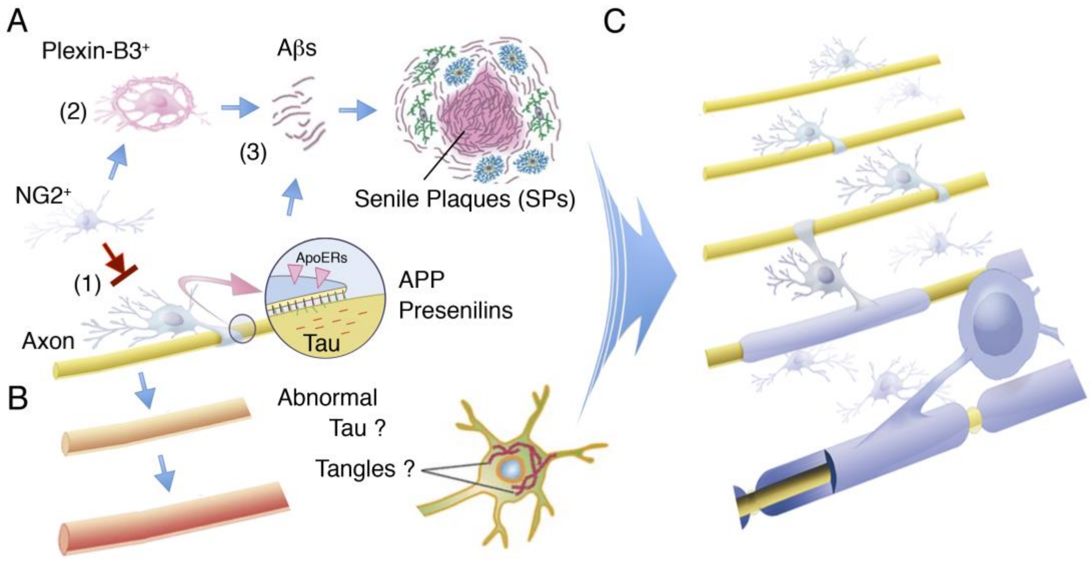


**Supplementary Fig. 12. Possible Alzheimer's disease pathology: implications from the present study.**

(**A**) A type of demyelination or dysmyelination occurs in the Alzheimer's disease cortex (1), resulting in defective plexin-B3+ aOPC differentiation (2) and extracellular Aβ accumulation (3). ApoERs: apolipoprotein E receptors expressing on a myelinating OL.
(**B**) Alzheimer's disease-type demyelination or dysmyelination may eventually promote neurological disabilities and probably formation of neurofibrillary tangles in neurons.
(**C**) Fine control of cortical myelination in aged brains might be an essentially requirement of effective Alzheimer's disease therapy.

| Brain area | Debris | Sup NG2% |
|---|---|---|
| Hippocampus | + | 93.5 ± 4.3 |
| Olfactory bulb | ++ | 87.7 ± 9.0 |
| Amygdala | ++ | 92.2 ± 5.0 |
| Striatum | ++ | 83.4 ± 13.7 |
| Cortex | +++ | 88.5 ± 6.5 |
| Medulla oblongata | ++++ | 78.4 ± 23.8 |
| Cerebellum | ++++ | 84.3 ± 3.4 |
| Thalamus and midbrain | +++++ | 85.4 ± 13.8 |
| Spinal Cord | +++++ | 91.7 ± 14.4 |

**Supplementary Table 1. Isolation and culture of aOPCs from various regions of adult rat brains.**

Using the Paxinos and Watson atlas (http://labs.gaidi.ca/rat-brain-atlas/) as a guide, the different brain regions were dissected from adult SD rats. aOPCs were cultured for 5 days, and the proportion of $NG2^{+}$ cells (Sup NG2%) was determined by the ratio (%, mean±SD) of immunolabeled cells to total cells (TO-PRO-$3^{+}$ cells). The amounts of debris were scored based on the impression of the observers.

| Antibody | Species | Source | Cat.# |
|---|---|---|---|
| NG2 | Rabbit | Millipore | AB5320 |
| NG2 | Mouse | Millipore | MAB5384 |
| PDGFR$\alpha$ | Rabbit | Santa Cruz | sc-338 |
| Olig2 | Rabbit | Millipore | AB9610 |
| O4 | Mouse | Millipore | MAB345 |
| GFAP | Mouse | Millipore | MAB3402 |
| GFAP | Rabbit | DAKO | Z0334 |
| Tuj1 | Mouse | Covance | MMS-435P |
| Neurofilament H | Rabbit | Millipore | AB1989 |
| GlyR$\alpha$2 (Glra2) | Goat | Santa Cruz | sc-17279 |
| Plexin B3 | Sheep | R&D | AF6879 |
| Notch 1 | Goat | Santa Cruz | sc-6015 |
| NICD | Rabbit | Cell Signaling | 4147 |
| GAPDH | Mouse | Santa Cruz | sc-32233 |
| APP(22C11) | Mouse | Millipore | MAB10424 |
| BACE1 | Rabbit | Calbiocam | 195111 |
| Presenilin 1 | Mouse | Millipore | MAB5232 |
| Human A$\beta$17-24 (4G8) | Mouse | Covance | SIG-39220 |
| Nicastrin | Rabbit | Signo Biological Inc | 11183-RP02 |
| Aph1(N-20) | Goat | Santa Cruz | sc-30240 |
| Rodent A$\beta$10-15 (M3.2) | Mouse | Covance | SIG-39155 |
| A$\beta$1-40 | Rabbit | IBL | 18580 |
| A$\beta$1-42 | Rabbit | IBL | 18582 |
| Tau [pS396] | Rabbit | BIOSOURCE | 44-752G |
| BrdU | Rat | serotec | OBT0030CX |
| Iba1 | Rabbit | Wako | 019-19741 |
| Iba1 | Rabbit | Wako | 061-20001 |
| PDGFR$\beta$ | Rabbit | Santa Cruz | sc-432 |
| Sox10 | Rabbit | Millipore | AB5727 |
| Nestin | Mouse | Millipore | MAB353 |
| CNPase | Mouse | Millipore | MAB326 |
| MBP | Mouse | Millipore | MAB384-1ML |
| CD68 | Mouse | DAKO | M0814 |

**Supplementary Table 2. List of antibodies used in the present study.**

A

| Case # | Age | Gender | Diagnosis | PMI | NFT | Amyloid | Weight |
|---|---|---|---|---|---|---|---|
| 1 | 63 | M | | 2h 41min | I | | 1515 |
| 2 | 75 | F | | 2h 6min | II | | 1280 |
| 3 | 78 | M | | 19h 56min | II | | 1270 |
| 4 | 68 | M | | 16h 2min | II | | 1460 |
| 5 | 78 | F | | 46h 56min | I | | 1200 |
| 6 | 80 | M | AD + DLB | 17h 11min | V | B | 1210 |
| 7 | 93 | M | AD | 19h 47min | VI | C | 1080 |
| 8 | 94 | M | AD | 81h 33min | V | B | 1290 |
| 9 | 96 | F | AD | 21h 43min | V | C | 1140 |
| 10 | 88 | F | AD | 76h 46min | V | C | 1190 |

B

| Case # | Age | Gender | Diagnosis | PMI | NFT Braak stage |
|---|---|---|---|---|---|
| 1 | 56 | M | | 8h | I/I |
| 2 | 60 | M | | 11h 32min | I/I |
| 3 | 87 | M | | 2h 32min | II/III |
| 4 | 57 | M | | 9h 20min | I/I |
| 5 | 87 | M | AD | 5h 50min | V/V |
| 6 | 66 | M | AD + DLB | 16h | V/V |
| 7 | 86 | F | AD | 7h 59min | IV/IV |
| 8 | 59 | F | AD + DLB | 3h | VI/VI |

**Supplementary Table 3. Demographic data of human brains.**
(**A**) Demographic data for the immunohistochemistry. Ten brains (5 Alzheimer's disease patients and 5 disease controls) from subjects autopsied at Nitobe Memorial Nakano General Hospital were included. Age: age at death (years), M: male, F: female, Diagnosis: Neuropathological diagnosis, DLB: diffuse Lewy Body disease, PMI: postmortem interval, NFT: Braak NFT (neurofibrillary tangle) stage, Amyloid: Braak amyloid stage, Weight: brain weight at autopsy (grams). (**B**) Demographic data for the WB analysis. Eight brains (4 Alzheimer's disease patients and 4 disease controls) from subjects autopsied at Aichi Medical University were included. Age: age at death (years), M: male, F: female, Diagnosis: Neuropathological diagnosis, DLB: diffuse Lewy Body disease, PMI: postmortem interval, NFT: Braak NFT (neurofibrillary tangle) stage, Amyloid: Braak amyloid stage, Weight: brain weight at autopsy (grams).
